# Parallel loss of sex in field populations of a brown alga sheds light on the mechanisms underlying the emergence of asexuality

**DOI:** 10.1101/2023.09.10.557039

**Authors:** Masakazu Hoshino, Guillaume Cossard, Fabian B. Haas, Emma I. Kane, Kazuhiro Kogame, Takahiro Jomori, Toshiyuki Wakimoto, Susana M. Coelho

## Abstract

Sexual reproduction is widespread among eukaryotes, but asexual lineages have repeatedly arisen from sexual ancestors across a wide range of taxa. Despite extensive research on the evolution of asexuality from sexual ancestors, the molecular changes underpinning the switch to asexual reproduction remain elusive, particularly in organisms with haploid sexual systems such as bryophytes, and red and brown algae in which males and females are haploid and multicellular. Here, we investigate independent events in which asexuality has emerged from sexual ancestor lineages in species of the brown algal *Scytosiphon*, we examine the proximate and evolutionary mechanisms involved, and test the importance of sexual conflict on gene expression changes following loss of sex. We find that individuals from asexual, female-only (‘Amazon’) populations lose their ability to produce and sex pheromone and, consequently, are unable to attract and fuse with male gametes, whereas they gain the ability to trigger parthenogenic (asexual) development from large, unfertilized eggs. This independent and convergent decline in pheromone production and optimization of asexual traits is accompanied by convergent changes in gene expression, including de-feminization and masculinization of the Amazon gamete transcriptomes. These data are consistent with the idea that decay of female functions, rather than relaxation of sexual antagonism, is the dominant force at play during the emergence of asexuality in haploid sexual systems. Moreover, we identify a locus on an autosomal protein-coding gene that is associated with the switch to asexuality. We propose that the sex chromosome, together with this autosomal locus, may underlie the switch to obligate asexuality in the Amazon populations.

## Introduction

The shift from sexual reproduction to asexual reproduction by parthenogenesis has occurred in several eukaryotic lineages (Neiman *et al*., 2014; Hojsgaard & Hörandl, 2019) but how the loss of sex affects the phenotype of the asexuals, the molecular changes involved in the switch and the genomic consequences of asexuality are unknown. Asexual lineages often occupy terminal nodes in phylogenies and are thought to be evolutionary dead-ends (Simon et al., 2003). According to theory, they are expected to have lower genetic diversity, reduced levels of adaptation, and generally lower fitness than their sexual relatives (Felsenstein, 1971; Keightley & Otto, 2006). However, empirical reports indicate that asexual populations frequently have similar diversity and rates of adaptation as their sexual relatives (see, for example, (Glemin & Galtier, 2012; Bast *et al*., 2018). It is still unclear, therefore, whether transitions to asexuality consistently bring the genomic changes that are predicted.

The extent to which the switch to asexuality is associated with large changes in gene expression is still debated. Asexual lineages may have a selective advantage over sexual lineages, as they are relieved from ‘intralocus sexual conflict’ (Abbott, 2011; Glemin & Galtier, 2012). Sexual conflict or sexual antagonism occurs when the two sexes have conflicting strategies to optimize their fitness for reproduction. If there is prevalent conflict between the two sexes concerning gene expression (for example, if the expression of a gene gives the male a fitness advantage but the female a fitness disadvantage), a shift to full asexuality would be expected to be accompanied by increases in expression of genes with primarily female functions (because they would be relieved from sexual antagonism), and decreases in expression of genes with primarily male functions. Yet, studies in stick insects found a masculinization of gene expression in asexuals (Parker *et al*., 2019; but see Huylmans *et al*., 2021). Thus, how the transition to asexuality impacts sexual conflict for optimal gene expression remains unclear. Moreover, the detailed molecular mechanisms underlying the shift to asexuality are largely unknown. Although hundreds of genes appear to be differentially expressed in sexuals versus asexuals (Liu *et al*., 2014; Parker *et al*., 2019; Huylmans *et al*., 2021), very few loci that are genetically associated with asexual states have been found (Dedryver *et al*., 2013; Ye *et al*., 2019).

Shifts to asexuality from sexual ancestors take place also in brown algae. These largely marine, multicellular eukaryotes evolved independently of animals and plants for billions of years; their last common ancestor was unicellular. This means that brown algae invented complex multicellular development independently of animals and plants. Their biology is largely underexplored, however, in recent years they have become key model organisms to study the evolution of sex determination, reproductive systems and complex life cycles (Coelho & Cock, 2020). Most brown algae alternate between multicellular male and female haploid gametophytes (the so called gametophyte generation) and a sporophyte (diploid) generation (‘haplo-diplontic’ life cycle) (Coelho & Cock, 2020; Heesch *et al*., 2021; Coelho & Umen, 2021; Cossard *et al*., 2022)(Figure 1A). Sex is determined at meiosis (not at fertilisation, as in XY and ZW systems) depending on whether haploid spores inherit a female (U) or a male (V) chromosome (Lipinska *et al*., 2015a; Umen & Coelho, 2019). Spores grow into multicellular female or male gametophytes, which at maturity produce female or male gametes (Figure 1A). In parallel with this sexual life cycle, brown algae may have an asexual life cycle in which unfused female (and occasionally male) gametes undergo parthenogenesis to develop into adult multicellular individuals (Peters *et al*., 2008; Bothwell *et al*., 2010a,b; Arun *et al*., 2013; Mignerot & Coelho, 2016; Mignerot *et al*., 2019).

**Figure 1.**
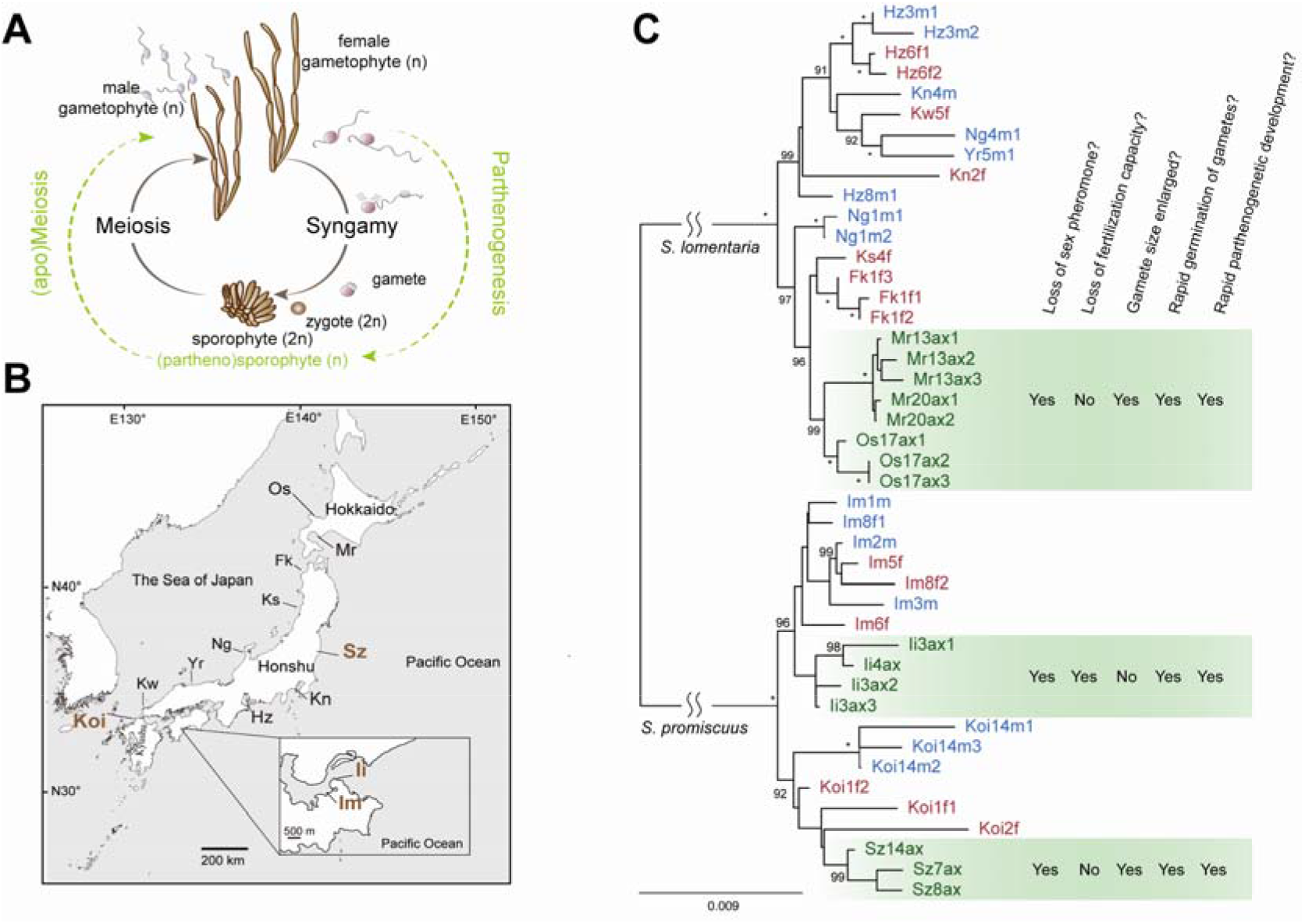
Amazon populations of Scytosiphon diverged from sexual ancestors < 1-2Mya. **A)** Haploid-diploid life cycle of Scytosiphon sp. Scytosiphon sp. has a heteromorphic life cycle, where generations of multicellular and macroscopic dioicous isomorphic gametophytes alternate with generations of microscopic multicellular discoid sporophytes. Scytosiphon sp. male and female gametes are almost indistinguishable in terms of size, with female gametes being slightly bigger than male gametes, but they show clear sexual dimorphism in behavior and physiology; female gametes settle on the substratum sooner than male gametes, and secrete a sex pheromone that attract male gametes (Fu et al., 2014). Note that unfused female gametes may enter a parthenogenetic cycle (in green) in absence of fertilisation, they develop into partheno-sporophytes which after apomeiosis reconstitute the gametophyte generation. Amazon populations use exclusively the parthenogenetic/asexual part of the life cycle. **B)** Geographical location of the different populations. Detailed location is given in Table S1. The four sampled populations of this study are highlighted in bold. **C)** Phylogenetic relationships between the studied populations. Maximum likelihood tree was based on concatenated DNA sequences of 53 single copy orthologues. Numbers on branches indicate bootstrap values from ML analysis. The asterisks indicate branches with full support (100). Only bootstrap values > 90 are shown. Summary of gamete behavior phenotypes scored in females, males and asexual (Amazon) females of different populations of S. promiscuus. Pheromone was measured either by olfaction in a blind test or by GC-MS (see methods). Attachment of male flagellum was measured by microscope observation (blind test). Gamete fusion was scored 24h after mixing male and female gametes by observation of zygote production (confirmed with presence of two eyespots), as in (Coelho et al., 2011).

Given their dramatic differences in terms of life cycle, the mechanisms of loss of sex (and transition to fully asexual reproduction) in haplo-diplontic organisms are expected to be fundamentally different from those described for animals, and, as comparative models, they provide an opportunity to explore the universality (or uniqueness) of this key biological transition. Previous work on the experimental model brown alga *Ectocarpus* found that parthenogenesis is controlled by the sex locus, and involves two additional autosomal loci (Bothwell *et al*., 2010a; Mignerot *et al*., 2019), highlighting the key role of the sex chromosome as a major regulator of asexual reproduction in this organism. However, the exact molecular mechanism underpinning the transitions to obligate parthenogenesis remains to be determined.

Transitions to obligate asexuality have been observed also in field populations of the brown alga *Scytosiphon* (Kitayama *et al*., 1992; Hoshino *et al*., 2019). Such asexual populations (referred to as ‘Amazon’ populations) and their closely related, sexual, ancestral populations provide a unique opportunity to study the transition from sexual reproduction to obligate asexuality in haplo-diplontic species. Moreover, the fact that transitions to asexual reproduction occur independently and repeatedly in these organisms allow us to investigate the conservation of the mechanisms underlying the switch to asexuality.

Here, we report the identification, and phenotypic and genomic characterization of asexual populations of a brown alga with haploid sex determination, and we contrast these populations with their sexual ancestors. We sequenced the gamete transcriptomes of several individuals from closely related pairs of asexual and ancestral sexual populations of two *Scytosiphon* species. This approach allowed us to test whether the pattern of gene expression consistently changes with the evolution of asexuality and whether there is a ‘feminization’ of gene expression in asexual females, consistent with a release from sexual antagonism. Contrary to expectations, we found that a female-specific trait (pheromone production) decays in Amazon populations, and there was a concomitant optimization of their capacity for asexual reproduction, by gamete parthenogenesis. These phenotypic modifications were accompanied by de-feminisation and masculinisation of gene expression. We identified a significant number of genes exhibiting convergent expression changes, greatly exceeding that expected by chance, including genes within the female sex-chromosome regions, which supports a role for sex-specific loci in the emergence of asexuality. A genome-wide comparative analysis of female and Amazon populations revealed one locus, encoding an EF-hand domain-containing protein that was fully associated with asexuality. This locus may play a role in signaling pathways related to the Amazon phenotype. Overall, by identifying changes occurring in multiple independent transitions to asexuality, our findings provide the first insights into the molecular basis of asexual reproduction in organisms with haploid sex determination and suggest that the evolutionary path to asexuality is highly constrained, likely requiring repeated changes in the same key genetic pathways.

## Results

### Amazon populations of *Scytosiphon* diverged repeatedly from sexual ancestors

We had previously identified natural populations of *S. lomentaria* composed exclusively of females (Hoshino *et al*., 2019). These Amazon populations appear to grow exclusively by asexual reproduction, in which female gametophytes develop following apomeiosis of parthenosporophytes formed upon parthenogenesis of unfertilized female gametes (Figure 1A). To investigate the prevalence of Amazon populations in the *Scytosiphon* genus, we sampled populations of the species *S. promiscuus* (closely related to *S. lomentaria*) at four sites (Koinoura, Koi; Shioyazaki, Sz; Inoshiri, Ii and Im) on the coast of Japan (Tables S1 and S2, Figure 1B), and determined their sex ratios by using sex markers (Lipinska *et al*., 2015a, 2017) and crossing experiments with reference strains. In total, 156 gametophytes were identified as *S. promiscuus* (Table S2). Two of the populations were sexual (Im and Koi), i.e., both male and females were present, and two populations were composed exclusively of females (Sz and Ii) (Table S2). These observations thus indicate that Amazon populations of *Scytosiphon* are relatively common and widespread, as are those of *S. lomentaria* (Hoshino *et al*., 2019).

To investigate whether the Amazon populations arose from ancestral sexual lineages across the distribution range, we examined the phylogenetic relationships between the *Scytosiphon* populations. We chose a subset of individuals from each population and sequenced RNA from gametes for each lineage (Table S1). RNAseq data was used to build a phylogenetic tree, which showed a clear separation of the *S. lomentaria* and *S. promiscuus* lineages, and revealed that Amazons emerged from their sexual ancestors repeatedly and independently in each population (Figure 1C). Based on the available data (see methods) we can estimate the divergence of asexual populations from sexual ancestors at < 1-2Ma (Figure S1).

### Decay of pheromone production and optimization of asexual traits in Amazon populations

Like most brown algae, *Scytosiphon* species are broadcast spawners: when the gametophytes reach maturity, female and male gametes are simultaneously released into the surrounding seawater. Female gametes rapidly settle and produce a pheromone, an aromatic C_11_H_16_ compound (Maier, 1995) that attracts male gametes. This compound can be detected by GC-MS or by an olfaction test (Maier and Müller 1986). Male gametes swim towards female gametes using their two flagella, they scan the membrane surface of the female gametes with their anterior flagella, then the gametes fuse to form a zygote, which develops into a (diploid) sporophyte (Figure 2A).

**Figure 2.**
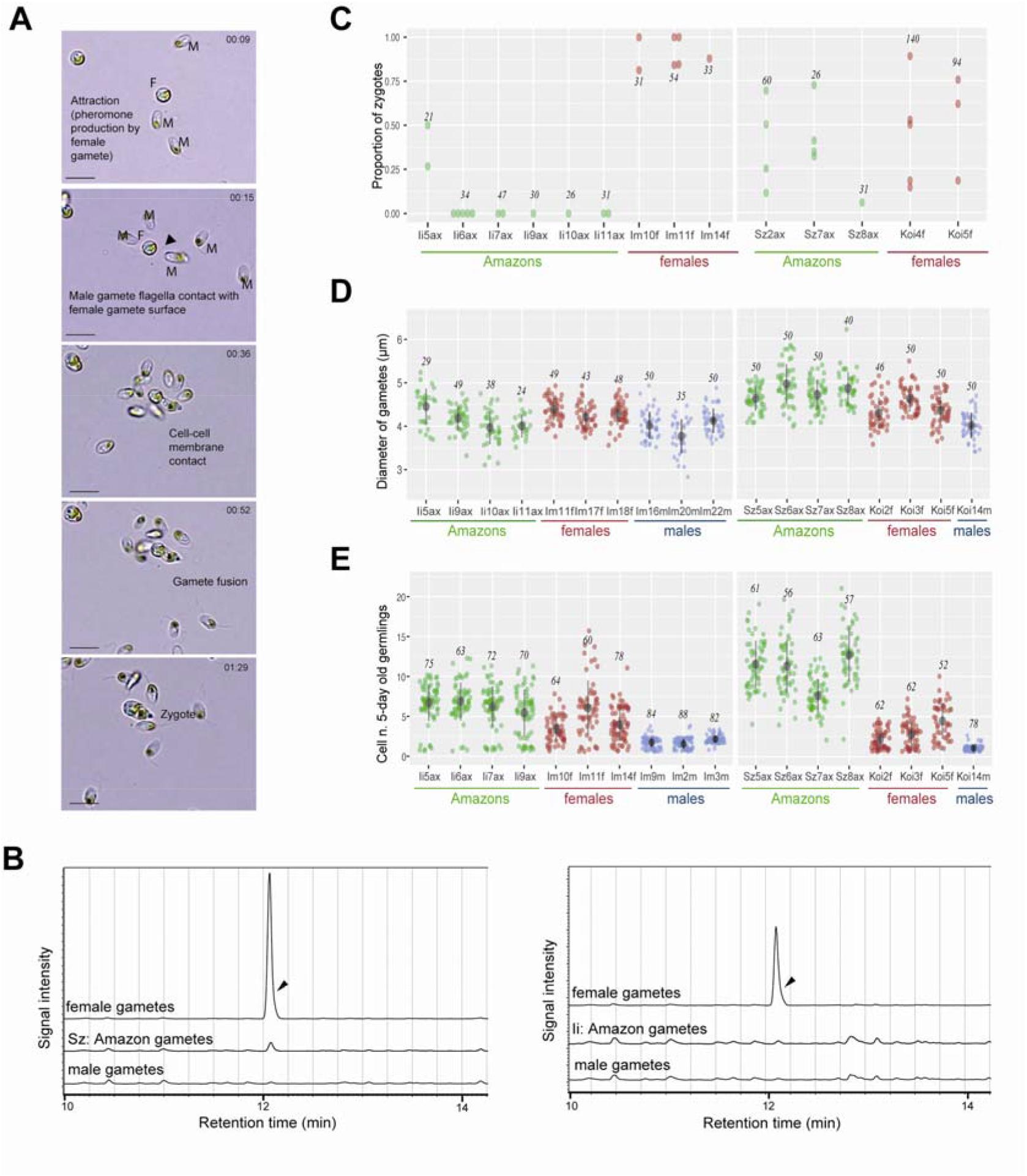
Decay of pheromone production and optimization of asexual traits in Amazon populations. **A)** Time lapse of fertilisation in a sexual population of S. promiscuus. Time (in seconds) is indicated in the top right of each photo. M: male gamete; F: female gamete. The first step is the attraction of male gametes toward female gametes via the production of the pheromone by the latter. Once male gametes are at the vicinity of the female gamete, they scan the surface of the female membrane using their flagella. The last step in the process is cell membrane contact and gamete fusion. **B)** Representative GC-MS analyses of the volatile compounds released from gametes: extracted ion chromatograms (MIC) = 91–92 m/z. The compound predicted as hormosirene (6-[(Z)-1-Butenyl]-1,4-cycloheptadiene) was detected with high intensity at around 12.1 min from female gametes of sexual populations (arrowheads). **C**) Fertilisation success of female gametes from sexual versus Amazon populations. We scored the proportion of zygotes arising in a controlled cross (see methods for details). Each point in the graph represents the fertilisation success of one independent cross. The number of scored germlings is indicated by the italicized number on each plot. **D)** Diameters of gametes in each line of S. promiscuus. The number of scored gametes is indicated by the italicized number on each plot, and each dot represent one gamete. The mean ± standard deviation is shown as a point range (black) in each plot. **E)** Parthenogenetic capacity of gametes, illustrated by the number of cells in developing 5-day old germlings in S. promiscuus. Female gametes may develop parthenogenetically into haploid partheno-sporophytes if they do not encounter a male gamete. Non-fused gametes from Amazon lines develop faster than un-fused gametes from sexual females. See also Table S3 and S4 for detailed statistical analysis.

We showed previously that gametes from *S. lomentaria* Amazon populations do not produce pheromone, although they still form zygotes with male gametes from sexual populations if gametes are incubated at high density in a small drop of seawater (Hoshino et al. 2021). We tested whether gametes from *S. promiscuus* Amazon populations, likewise, were unable to produce the sexual pheromone. GC-MS analysis detected an C_11_H_16_ compound in female gametes from sexual populations of *S. promiscuus* but failed to detect this pheromone or other C_11_ hydrocarbons in samples from any of the Amazon strains of this species (Figure 2C, Table S3). This suggests that Amazon female gametes of both species do not attract male gametes by producing pheromone.

To test whether gametes from *S. promiscuus* Amazon populations can fuse with male gametes from sexual populations in laboratory conditions, we recorded the behavior of Amazon gametes when the male gametes were present at high density to ‘force’ the encounter between them. In these conditions, most Amazon Ii populations did not form zygotes (Figure 2C), indicating little or no fertilization. In contrast, related sexual populations (Im) produced a large proportion of zygotes. We also noticed that male gamete flagella did attach to the membrane of the Amazon Ii gametes, but this recognition resulted in gamete fusion only in one sample, Ii5ax (Figure 2C). By contrast, Amazon Sz gametes formed as many zygotes as did the relates sexual (Koi) female gametes in the conditions of high male gamete density (Figure 2C, Tables S3 and S4, Generalized Linear Mixed Model (GLMM) model), indicating effective fertilization. The ability of Amazon gametes to recognize and fuse with the male gamete membrane was unaffected in all populations except in population Ii, in which, despite the high density of male gametes, no gamete fusion occurred (Figure 2C). This suggests that the genetically female Amazon Ii population produces ‘zoids’ that cannot fuse with gametes of the opposite sex; it is fully asexual. The zoids from the other Amazon populations are affected only in their capacity to produce pheromone.

We performed a morphometric analysis of the gametes of Amazon and sexual populations to look for differences between them. *S. promiscuus* Amazon gametes tended to be bigger than female gametes from sexual populations, significantly so for population Sz (Figure 2D, Table S4, GLMM model). Moreover, the parthenogenetic capacity of the Amazon gametes, determined by germination rate of unfused gametes after 24h, was significantly greater than that of their closest sexual populations (GLMM model, Table S4). Amazon parthenogenetic germlings were morphologically more developed than their sexual counterparts after five days in culture, as indicated by the number of cells they contained (Figure 2E, Table S4, GLMM model).

Our observations suggest that genetically female Amazon populations of both *S. promiscuus* and *S. lomentaria* do not produce a sex pheromone to attract male gametes. Despite the convergent decay of this sexual trait, Amazon gametes of both species retain their ability to recognize male gametes and fuse with them, except for Amazon gametes from the Ii population. This indicates that they have not become fully asexual. Concomitant with loss of pheromone production, Amazon gametes undergo parthenogenesis more rapidly and grow faster than female sexual gametes, therefore, they appear to be optimized for asexual reproduction.

### Transcriptome landscape in males, females and Amazons

To determine whether the phenotypic changes in the Amazon gametes were accompanied by modifications in their transcriptomic landscape when compared with male and female gametes, we used RNA-seq to measure transcript abundance in male and female gametes from the sexual populations and in Amazon gametes (Tables S5 and S6). To map the transcriptomes, we generated high quality whole genome assemblies for both *S. lomentaria* and *S. promiscuus* by using a combination long-and short-read sequencing (Tables S5 and S6).

The number of genes expressed in the gametes was 11,378 in *S. lomentaria* (57% of the genome) and 12,809 in *S. promiscuus* (61% of the genome; Tables S5 and S6). In principal component analysis (PCA) using all samples from female sexual and Amazon populations, samples clustered by species and by population; the females and Amazons did not form distinct clusters (Figure 3A). It appears, therefore, that there is no extensive modification of transcription when all genes are compared between sexual females and Amazon, probably reflecting their relatively recent split. However, when PCA was performed separately for each species, and male, female and Amazon samples were included, the Amazon populations clustered together (Figure 3B). This implies that the transcriptomes of Amazon populations are more closely related to each other than to their phylogenetically closest female populations.

**Figure 3.**
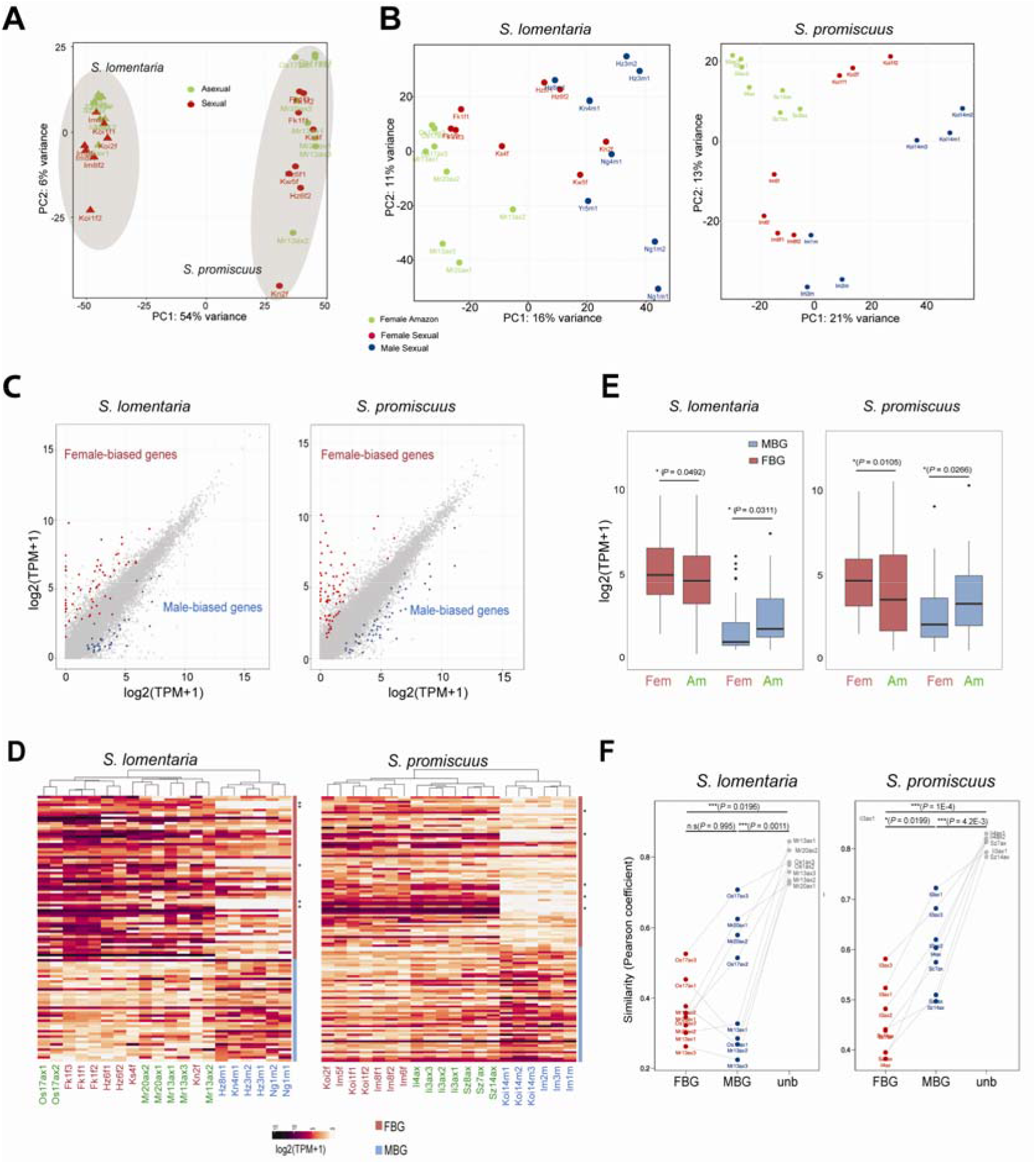
Defeminization and masculinisation of gene expression associated with asexuality. A) PCA plot of female and Amazon RNA-seq samples for both S. promiscuus (Sprom) and S. lomentaria (Slom), using SCOs. B) PCA based on all expressed genes per species. Sexual female samples are represented by a dot, Amazon (asexual) female samples by a triangle. C) Comparison of gene expression levels, in log2(TPML+L1), between males and females in S. lomentaria and S. promiscuus. Dark pink points represent female-biased genes, blue points male-biased genes. Grey points represent unbiased genes. D) Hierarchical clustering and heatmap of gene expression (in log2(TPM+1)) for all the SBG in each species (ComplexHeatmap package, R). The dendogram was constructed using hierarchical clustering based on Euclidean distances (defaults parameters in ComplexHeatmap). Asterisks indicate genes that are female-linked (inside the U-SDR), therefore absent from male samples. Sexual females, males and Amazons are marked in red, blue and green, respectively. E) Comparison of gene expression levels in log2(TPML+L1), using SCO gene sets, during transition to asexuality in S. lomentaria and S. promiscuus. The boxes represent the interquartile ranges (25th and 75th percentiles) of the data, the lines inside the boxes represent the medians, and the whiskers represent the largest/smallest values within 1.5 times the interquartile range above and below the 75th and 25th percentiles, respectively. The statistical tests are Mann-Whitney ranked tests, p-values displayed in parenthesis. F) Comparison of similarity index values (Pearson coefficients) between expression profiles (in log2(TPM+1)) of Amazon and sexual females for unbiased, female, and male-biased genes. Significant differences of coefficients between sex-biased genes are indicated directly on the plot.

Our phenotypic analyses, described above, revealed that Amazon gametes have lost a female-specific trait, the ability to produce pheromone. To examine whether sex-biased gene expression underlies this phenotypic ‘de-feminization’, we analyzed the RNA-seq data with DEseq2 to identify gene sets that are expressed differentially in male versus females in the sexual strains (i. e., sex-biased genes). The analysis considered only genes that displayed at least a two-fold change (FC) in relative expression between the sexes (Table S6; FCD>D2, *P*_adj_D<D0.05). Less than 1% of the expressed genes in *S. lomentaria* and *S. promiscuus* gametes were sex-biased genes (Figure 3C, Table S6). This is consistent with the overall low level of sexual dimorphism in brown algae (Luthringer *et al*., 2015). We found similar proportions of male-biased genes and female-biased genes in both species (Tables S6–S8, Figure 3C). Functional enrichment analysis of sex-biased genes highlighted functions related to catabolism (peptidase activity) specifically in female-biased genes in both species (Figure S3, Table S9). In other species with UV sex chromosomes, sex-biased genes evolve faster than unbiased genes (Lipinska *et al*., 2015b). We tested if this was the case for *Scytosiphon* sp. by comparing the evolutionary rates (dN/dS) of 4,747 expressed single-copy orthologs (SCO) in *S. lomentaria* and *S. promiscuus* that we identified by using the OrthoFinder algorithm (Table S10). For each species, we compared the evolutionary rates of SCO that have sex-biased expression with those of SCO whose expression is unbiased. The dN/dS values of female-biased genes were significantly greater than those of the unbiased genes, indicating a faster evolutionary rate, whereas we saw no evidence for faster evolution of male-biased genes (Figure S3).

To investigate how the expression of these sex-biased genes was modified in Amazon populations, we analysed the RNA-seq data for all the sex-biased genes in each species by hierarchical clustering in a heatmap. Overall, the sex-biased gene transcriptomes of females and Amazon samples in both species were more similar to each other than they were to male samples; however, the Amazon samples clustered together and formed a separate cluster from female samples, particularly in *S. promiscuus* (Figure 3D). This suggests that sex-biased gene expression in Amazons differs in some ways from that of a fully functional female.

The median level of female-biased gene expression [measured as log_2_(TPM + 1)] in Amazon females was significantly lower than that of sexual females (Wilcoxon test, *P*=0.049 and *P*=0.010 in *S. lomentaria* and *S. promiscuus* respectively; Figure 3E), suggesting these genes are transcriptionally ‘de-feminised’ in Amazons. By contrast, male-biased genes were all expressed at a higher level, or a similar level, in Amazons when compared to sexual females (Wilcoxon test, *P* =0.003 and *P*=0.02 in *S. lomentaria* and *S. promiscuus*, respectively; Figure 3E, Figure S3), suggesting that Amazon gametes are transcriptionally ‘masculinized’. As expected, there was no overall change in sex-biased gene expression when two sexual lineages were compared, (Figure S4). Thus, we conclude that there was a *bone fide* de-feminisation and masculinisation of gene expression upon transition to asexual reproduction.

Together, these findings indicate that transition to asexual reproduction in Amazons involves systematic changes in sex-biased gene expression in gametes, leading to de-feminisation and masculinization of gene expression, when compared with females.

To characterise further the changes in expression of sex-biased genes upon transition to asexuality, we calculated the mean expression of female-biased, male-biased and unbiased genes in females and Amazons of *S. lomentaria* and *S. promiscuus* from all the population samples and correlated expression of genes in the asexual samples with expression of the orthologous genes in the female sample by calculating the Pearson correlation coefficients. A correlation coefficient of 1 indicates a very similar level of expression, whereas correlation coefficients < 1 indicate increasing differences in expression compared to ancestral sexuals. We found that the levels of unbiased gene expression in asexual samples were very similar to those in the sexual samples, whereas female-biased and male-biased gene expression in asexual samples were very different to those in the sexual samples (Figure 3F). The effect was more marked for the female-biased genes than for the male-biased genes in most populations. In other words, the transition from female to asexual involves major changes in transcription, mainly at the level of sex-biased genes and mostly so in female-biased genes.

### Conservation of the transcriptomic changes associated with the switch to asexuality

In animals, the events that result in emergence of asexuality from sexual ancestors involves convergent changes in gene expression, i.e., in each independent transition to asexuality the same genes show similar changes in their pattern of expression (Parker *et al*., 2019; Huylmans *et al*., 2021). We thus analysed whether the decay in pheromone production that we observed in the various Amazon populations was accompanied by convergent modifications of their transcriptomes upon each transition to asexuality. The analysis compared the set of SCO common to *S. lomentaria* and *S. promiscuus* (Table S10). Of all expressed SCO, 6.7% (i.e., 320 of the 4776) exhibited convergent expression, i.e., they either up-regulated or down-regulated their expression in the same way in each of the independent transitions to asexuality (Tables S10 and S12; see for example the SCO in Figure 4A,). This proportion was significantly greater than what would be expected by chance (determined by using permutation tests, *P*=0.0419, 10,000 permutations). Functional analysis of these convergently expressed genes identified functions related to oxidative metabolism and ion transport (Figure 4B, Table S13). These convergent genes displayed no differences in their evolutionary rates compared to non-convergent genes (mean dN/dS = 0.195 in genes with convergent expression shifts; mean dN/dS = 0.191 genes without convergent expression shifts; *P* = 0.568, paired Mann-Whitney U-test). Thus, the transition to asexuality involves a small but significant number of genes that consistently change their expression patterns. Yet, these genes show no difference in their evolutionary rates suggesting their sequences are not under selective pressure.

**Figure 4.**
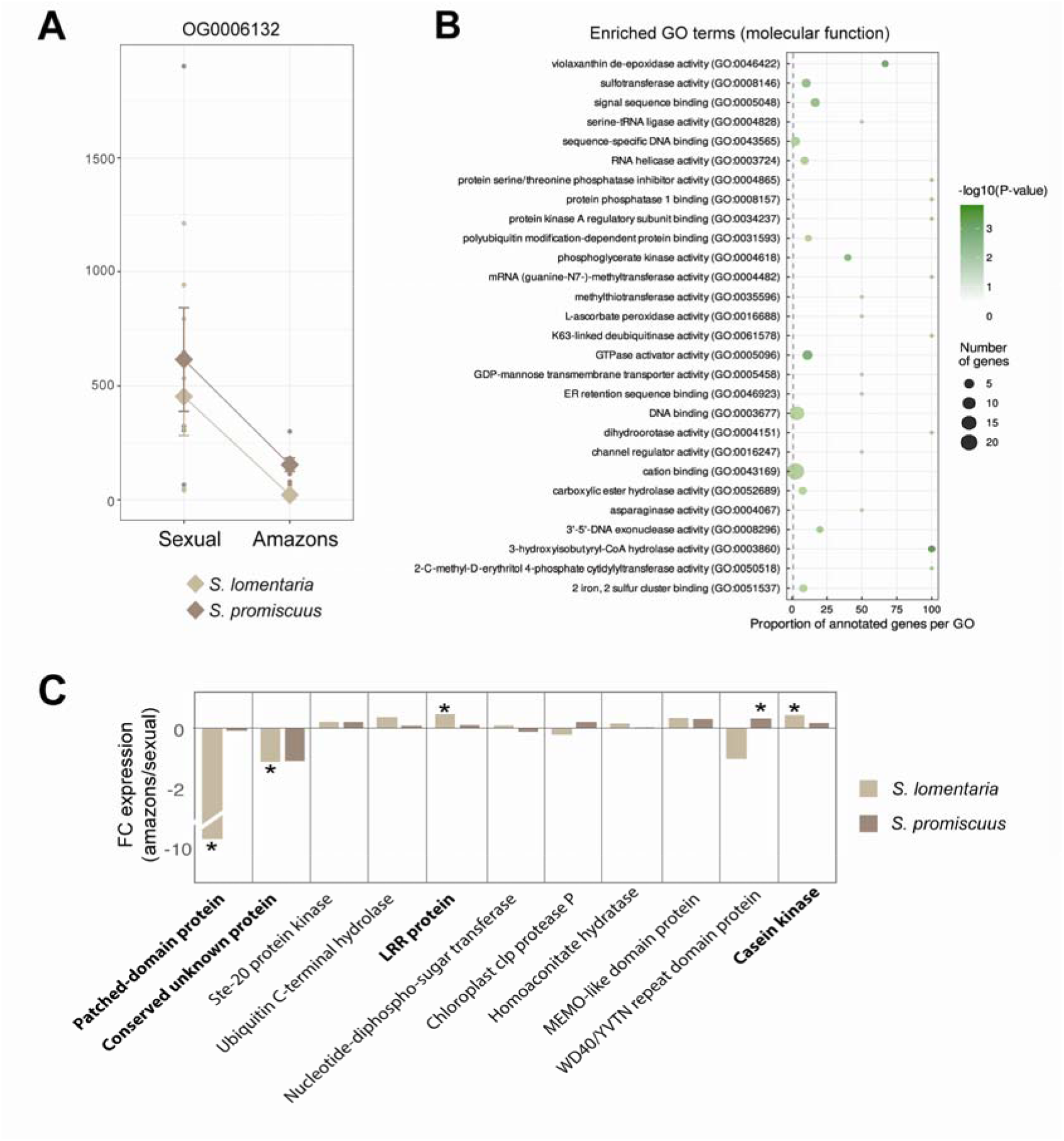
Convergent gene expression changes during transition to asexuality. **A**) Example of a single copy ortholog (SCO) (OG0006132) showing convergent gene expression changes in both species during transition to asexuality. The plot of normalized counts was generated using the PlotCount function in Deseq2 package. See also Table S10 for details of expression of each SCO showing convergent expression. **B**) Gene ontology (GO) enrichment (molecular function) in genes that exhibit convergent expression changes during the transition to asexual reproduction. Note that terms related to cellular process and cellular component are presented in Table S9. **C**) Changes in expression level (fold change, FC) of female U-linked genes in Amazon (asexual) populations compared to ancestral sexual populations. Asterisks indicate significant differences (Wilcoxon test, p<0.05). Protein-coding genes which have similar expression changes in the two species during the transition to asexuality are highlighted in bold.

The lack of pheromone production in Amazons appears to be fixed in these populations, consistent with previous work suggesting that the lack of pheromone production is inherited in F1 hybrid females of an Amazon and a male in *S. lomentaria* (Hoshino et al., 2019). This raises the possibility that the female sex-determining region of the U chromosome (U-SDR) is involved in the pheromone pathway. If so, we might expect to find differences in expression of U-SDR genes in females and Amazons of both *S. lomentaria* and *S. promiscuus*, where transition to asexuality occurred independently. SDR genes have been described for *S. lomentaria* based on their homology with those of the brown alga *Ectocarpus* (Lipinska et al., 2017); we verified that these genes were also specific to the genome of female *S. promiscuus* and that they were expressed specifically in females.

We compared the expression levels of U-SDR genes in females and Amazons. When all U-SDR genes were analyzed together, we found no significant difference in their expression between females and Amazons of either species (Figure S6). Nonetheless, a few U-linked genes were consistently up-or down-regulated in the Amazons of both species (Figure 4C). Notably, a U-specific gene predicted to encode a transmembrane protein containing a Patched domain, similar to the Patched membrane receptor of the Hedgehog signalling pathway, was significantly down-regulated in Amazon populations of *S*. *lomentaria* when compare with females and in at least one of the Amazon lineages of *S. promiscuus* (Table S11, Figure S6). This gene is part of the ancestral group of genes that are thought to have been present in the ancestral female sex locus of brown algae (Lipinska *et al*., 2017).

Together, these data indicate that whereas expression of most U-SDR genes was similar in Amazons and females, certain genes in this genomic region were differentially expressed in both *Scytosiphon* species, consistent with the transition to asexuality.

### Reproductive mode and selection efficacy

Loss of recombination in asexual organism is predicted to result in changes in the number of neutral polymorphisms segregating in the population and reduced effectiveness of selection (reviewed in (Hartfield, 2016). To investigate whether the Amazon populations of *Scytosiphon* species had fewer polymorphisms than sexual populations, as has been shown in asexual diploid organisms (Neiman *et al*., 2010; Hollister *et al*., 2015; Bast *et al*., 2018) we compared the proportions of genes containing single nucleotide polymorphisms (SNPs). We found a greater proportion of genes containing SNPs in sexual populations of both *S. promiscuus* (P < 2.2 x10^-16^) and *S. lomentaria* (P = 6.32 x10^-16^) when compared with Amazons. We used a generalized linear model with binomial distribution to analyze the proportion of variable sites among genes that contained SNPs. Sexual populations had a greater proportion of variable sites than had Amazon populations, both in *S. promiscuus* (P < 2.2×10^-16^) and in *S. lomentaria* (P < 2.2 x10^-16^) (Figure 5A-B).

**Figure 5.**
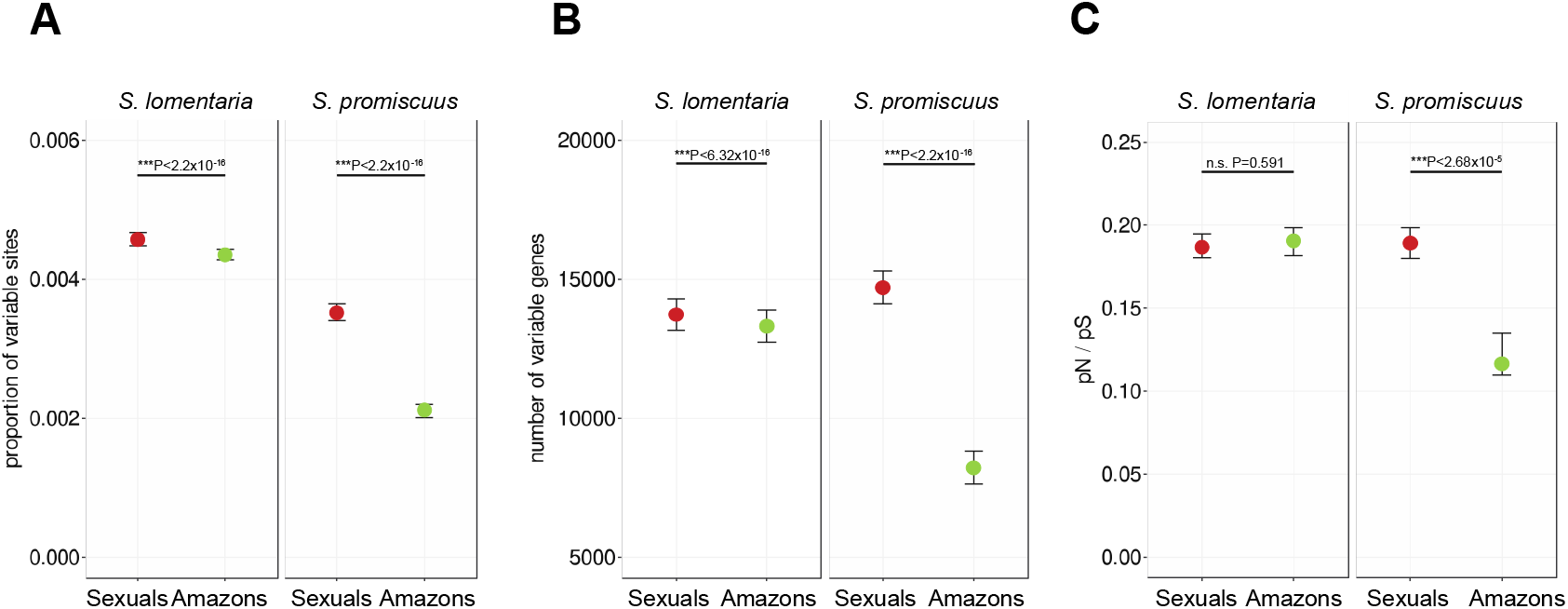
Selection efficacy in sexual versus Amazon populations of S. lomentaria and S. promiscuus populations. **A**) Proportion of variable sites in sexual and Amazon populations. **B**) Number of genes presenting variable sites. **C**) pN/pS [representing rates of non-synonymous (pN) and synonymous (pS) polymorphism] in sexual versus Amazon populations.

To test the prediction that the absence of sex and recombination in Amazon lineages results in reduced effectiveness of selection, we looked for evidence of decreased purifying selection. Purifying selection removes deleterious mutations from a population, thus decreased purifying selection results in accumulation of non-synonymous mutations in protein-coding genes, which are mostly deleterious. The ratio pN/pS is predicted to be higher in asexual species when compared with sexual species (Normark & Moran, 2000). Note that because gene expression is likely to affect pN/pS (Drummond et al., 2005; Park et al., 2013), we included it [in log_2_(TPM+1)] in our models as a dependent variable. Also, to infer the effect of asexuality on pN, we conducted likelihood ratio tests with a reduced model without reproductive mode (sexual, asexual). We found no statistically difference between the mean values of pN/pS in sexual and Amazon populations of *S. lomentaria* (GLMM P = 0.719), whereas the mean pN/pS in sexual populations of *S. promiscuus* was much greater than that of Amazon populations (GLMM, P < 2.2 x10^-16^)(Figure 5C).

We performed the same calculations for the subset of genes in each species that belong to a SCO, as we expect these long-time conserved genes to be maintained under strong purifying selection. The trends of pN/pS in SCO were similar to those inferred from all genes within species, i.e., the mean value of pN/pS in sexual lineages was no different (in *S. lomentaria*) or greater (in *S*. *promiscuus*) than that of Amazons in (Table S14).

### An EF-hand domain-containing protein associated with the Amazon phenotype

To investigate if any changes at the genomic level were consistently associated with the transition to asexuality in *S. lomentaria* and *S. promiscuus* females, we searched for genetic variants associated specifically with asexuality in the transcriptomes of females and Amazons. In S. *promiscuus*, we identified 16 transcripts in *S. promiscuus* containing non-synonymous mutations that were fully associated with the Amazon phenotype (Table S14). Remarkable among these 16 variants, one had a missense mutation that was also fully associated with the Amazon phenotype in *S. lomentaria*. This transcript encodes an EF-hand-containing protein. In *S. lomentaria* the mutation results in replacement of a glycine residue by a tryptophan (Gly511Trp), whereas in *S. promiscuus* it results in replacement of a threonine residue by an arginine (Thr516Arg)(Figure 6A; Table S15). The mutations occurred in the same exon in both species and correspond to a similar position in the primary sequence. Based on the positions of these species on the phylogenetic tree (see Figure 1B), we conclude that these mutations occurred independently.

**Figure 6.**
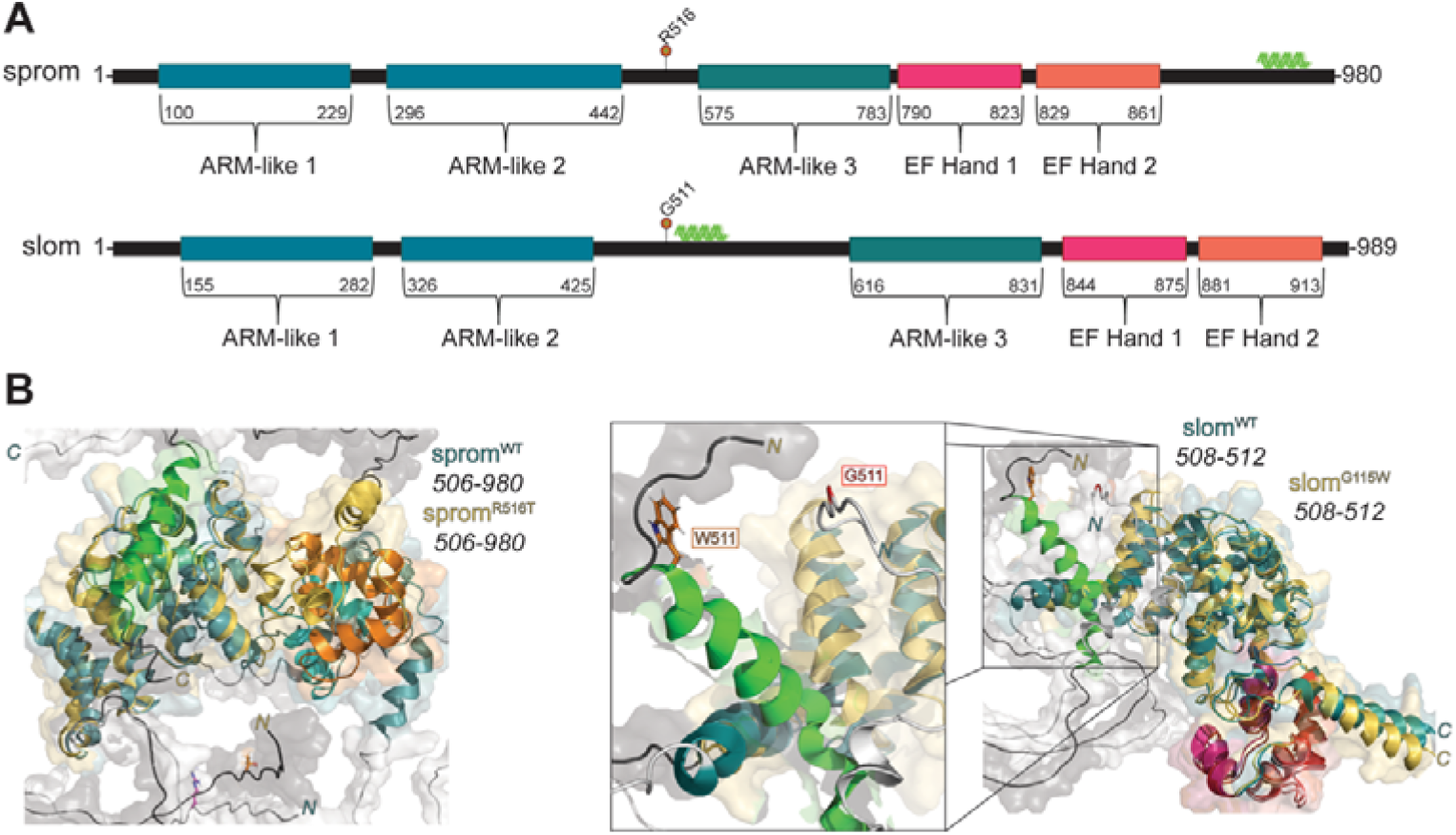
Domain architecture and models of EF hand-ARM homologs from S. lomentaria and S. promiscuus. **A)** Protein domain architecture of both protein-coding genes from S. lomentaria and S. promiscuus. Both orthologs contain three ARM-like repeat domains (dark blue) that encapsulate large, unfolded loops, as well as two C-terminal EF-Hand motifs (magenta/orange). Both variant sites are highlighted (red lollipop) and precede the third ARM-like domain. The additional α-helix predicted with each mutant protein is highlighted (light green). **B)** Generated models of the wild-type (sexual) and variant proteins from each species. The Thr516Arg substitution is distal to the EF-hand motifs (left) and are predicted to have little impact on the structural integrity. While both variants have an additional α-helix that is within proximity to the C-terminal ARM domains, only the Gly115Trp mutation within the S. lomentaria variant initiates the newly acquired α-helix (right) following the mutation site. ARM, Armadillo-like repeat domain; C2, EF Hand, Eps15 Homology Hand motif.

We used molecular modelling to predict the structures of these proteins and the effects of the mutations on protein structure and function. The primary structures of the two proteins are highly conserved between the two species (87.2% identity); they contain three armadillo-like repeat (ARM-like) domains at their N-termini and two EF-hand domains at their C-termini based (Figure 6A). Armadillo-repeat domains are typically involved in protein-protein interactions and intracellular signaling. Predictions of the folded, three-dimensional structures of the wild-type (sexual) and variant (Amazon) forms of each ortholog made by using AlphaFold2 indicate that the structures of the EF-hand domains are little affected by the mutations (Figure 6B), thus the calcium-binding functions of both Amazon proteins probably remain functional, at least to some extent (Figure 6B). The additional alpha helix induced by the mutations in the variants of the Amazons is expected to lead to altered protein-protein interactions.

## Discussion

Here, we have studied field populations of two species of the brown alga *Scytosiphon* to determine the phenotypic and molecular modifications that are associated with the transitions to asexuality in organisms with a haploid-diploid life cycle, where females and males are multicellular haploid organisms.

### Emergence of obligate asexuality is associated with loss of pheromone production and optimization of parthenogenesis

Asexual populations, which reproduce exclusively by female gamete undergoing parthenogenesis, have emerged repeatedly from sexual ancestors and appear to be frequent in the genus *Scytosiphon*. Female-specific pheromone production is lost consistently during the switch to asexuality, and this loss is likely to be fixed in these populations because F1 females have been shown to be incapable of pheromone production (Hoshino *et al*., 2018). The fact that F1 females do not produce pheromone is consistent with a major pheromone locus being inside the U-SDR, that is transmitted to all daughters. Substantial further work will be needed to identify precisely the complex genetic basis of pheromone production. Interestingly, the loss of pheromone production is conspicuous also in animals, during their transition to obligate parthenogenesis (Parker *et al*., 2019) supporting the idea this is a costly trait that is rapidly dispensable in the absence of males.

In all but one population of Amazon we examined, the gametes were still able to recognize and fuse with male gametes of the same species, despite having lost the capacity to attract male gametes by producing pheromone. This observation suggests that the transitions to asexuality have occurred relatively recently and have not led to full loss of sexual capacity. Accordingly, our estimations indicate that the Amazons populations emerged <1-2 Mya.

Concurrent with the loss of pheromone production, Amazon female gametes are often larger than the sexual female gametes. This suggests that after the evolution of obligatory parthenogenesis, the female gamete becomes larger to compensate for the lack of resources provided by the male gamete. In addition, Amazon female gametes rapidly engage in parthenogenetic development. Together, these observations indicate that Amazon female gametes are specialized for asexual reproduction by parthenogenetic development of unfertilized gametes. In the closely related brown alga *Ectocarpus* which also has a haplo-diplontic life cycle, gamete parthenogenesis is controlled by a major quantitative trait locus located in the U-SDR. It has been suggested that this trait may be subject to both sexual selection and generation/ploidally-antagonistic selection (Mignerot *et al*., 2019). In other words, parthenogenesis may be advantageous in situations where there is no pheromone or when males are absent, whereas producing the pheromone is costly and is only advantageous in presence of males. If parthenogenesis is not costly, then it can be maintained more easily than pheromone production. This mechanism would be consistent with a trade-off (Stearns, 1989; Johnston *et al*., 2013) between the haploid and diploid stages of the life cycle, where distinct parthenogenesis alleles have opposing effects on sexual and asexual reproduction and may help maintain genetic variation (Mignerot *et al*., 2019). In the case of Amazon populations, the allele that confers fast and efficient parthenogenesis would be favored and would rapidly spread in the population because it would ensure higher survival in situations where gamete encounter is limiting.

### Origin of Amazon populations

In animals and land plants, interspecific hybridization and polyploidy are major triggers for parthenogenesis (Simon *et al*., 2003; Kearney, 2005; Neiman *et al*., 2014). In the brown alga *S*. *lomentaria*, however, phylogenetic analyses based on mitochondrial and nuclear markers have found no evidence for interspecific hybridization (Hoshino *et al*., 2021). At least two scenarios are possible to explain the origin of the Amazon populations. In the first scenario, a mutation leading to loss of pheromone production triggers the loss of sex (i.e., ‘spontaneous origin’, (Simon *et al*., 2003). Because female gametes are capable of parthenogenesis, then the female-parthenogenetic population expands whereas males would decay because they cannot undergo efficient parthenogenesis (Hoshino & Kogame, 2019). In the second scenario, males are less tolerant to a change in environmental conditions (e.g. water temperature, (Hoshino & Kogame, 2019; Hoshino *et al*., 2021), so they become less frequent, which favours females that rapidly reproduce by parthenogenesis. As pheromone production is presumably costly, this trait would then be lost. In this scenrario the loss of pheromone is secondary to the loss of sex.

Interestingly, our Amazon populations did not undergo the mutational meltdown that is predicted to arise in parthenogenetic populations due to the absence of recombination. We did not see the strongly reduced efficacy of purifying selection, as observed in stick insects (Bast et al, 2018). Ancient asexuality would allow the purging of deleterious mutations (Brandt *et al*., 2017), but our phylogenies do not support this hypothesis because the emergence of Amazons appears to be relatively recent. Because Amazons spend their entire life cycle in the haploid stage (see Figure 1), it is likely that haploid purifying selection efficiently purges deleterious mutations (Szövényi *et al*., 2014; Brandt *et al*., 2017). Moreover, one intriguing possibility is that several different females from an ancestral sexual population were at the origin of the Amazon populations and this would have contributed to have a pool of diversity in the Amazons.

### Defeminization of gene expression driven by decay of female traits?

Males and females have distinct phenotypes, despite sharing most of their genome. A possible resolution of this apparent conflict is by differential gene expression, whereby genes are expressed at different levels in each sex (Connallon & Knowles, 2005). This differential gene expression, however, may lead to conflicts between the optimal expression of genes in males and females. Our data allowed us to test the hypothesis that gene expression in (haploid) females is constrained from evolving to its optimum level due to sexually antagonistic selection on males, by examining changes in sex-biased gene expression in the Amazon populations, which do not produce males. We expected the transcriptome of Amazons to be feminized, as they no longer experience anymore sexual conflict, thus gene expression could reach its female-optimum. However, as in animals (Parker *et al*., 2019; Huylmans *et al*., 2021), gene expression was defeminized, simultaneous with loss of the female-specific trait of pheromone production. We propose that the changes in sex-biased gene expression during the switch to asexuality in this organism are thus mainly related to the loss of (costly) female traits and consequent changes in trait optima, and not necessarily to resolution of sexual conflict. Since sexually dimorphic traits are thought to be a product largely of sex-biased gene expression (Mank, 2017), a link between reduced female sexual traits and reduced female-biased gene expression is a plausible explanation for the decreased expression of female-biased genes we observe and to the dramatic change in female-biased gene pattern of expression (low Pearson similarity index).

### Transcriptomes of Amazon populations converge

The independent, convergent decay in pheromone production that we observed in the various Amazon populations was accompanied by convergent modifications of their transcriptomes upon each transition to asexuality. The transcriptomes of Amazons were more closely related to each other than to those of their phylogenetically closest sexual population, suggesting similar regulatory networks are recruited repeatedly and independently during the transition to asexuality. The convergent gene expression changes that were associated with the transition to asexuality did not, however, appear to be driven by selection. It is possible that rapid engagement in parthenogenesis to develop the partheno-sporophyte generation (Figure 1) explains the convergence in transcriptomes. Interestingly, convergently recruited genes in Amazon populations have putative functions similar to those enriched in the sporophyte generation of *S. lomentaria* (Lipinska et al., 2019). Considering that Amazons are optimized for parthenogenesis and rapid (partheno)sporophyte development (see above), it is therefore likely that their changes in gene expression reflect a switch in life cycle strategy.

The U-specific region harbors several genes that converge in terms of their changes of expression during the switch to asexuality. Whilst phenotypic sexual dimorphism is not necessarily caused by genes within the non-recombining sex determining region, genes within the U-SDR could eventually be involved in the phenotypic changes that underlie the transition to asexual reproduction. Interestingly, a major locus controlling parthenogenesis is linked to the SDR in a closely related brown alga (*Ectocarpus* sp.) and it is likely that pheromone production may also be controlled by the sex locus. It is impossible, however, to determine whether the convergent changes in expression of SDR genes are a cause or effect of the asexuality; SDR genes that are differentially regulated in Amazons may be involved in triggering asexuality or may simply be a consequence of this developmental switch.

Remarkably, the transition to asexuality was genetically associated with a change in the protein coding sequence of a gene encoding a putative Armadillo domain-and EF-hand domain containing protein. EF-hand domains bind Ca^2+^, thus this protein is likely involved in intracellular Ca^2+^ signaling. Reverse genetic tools are not yet available for *Scytosiphon* therefore the causal relationship cannot currently be established between this protein and asexuality, but our data provides compelling evidence that sequence variation in this gene is associated with the Amazon phenotype. Ca^2+^ plays an important role in signaling pathways in brown algae (Coelho *et al*., 2002, 2008) and in pheromone signaling pathways in other eukaryotes (Ma & Roelofs, 1995) and Armadillo domains have well-characterized roles in protein-protein interactions and membrane-binding (Tewari *et al*., 2010). It is possible that a membrane binding ability of the Amazon variant protein is compromised, or a protein-protein interaction is disrupted, that is essential for Ca^2+^-dependent pathways during sexual reproduction and/or pheromone production.

## Methods

### Phenotyping

We focused on four gamete phenotypic traits: fertilization rate when crossed with gametes of the opposite sex, sex pheromone production, gamete size, and capacity for parthenogenetic development. These phenotypic traits were examined in female-only, asexual populations, Ii and Sz, and they were contrasted with the closest related sexual (ancestral) populations (i.e., Ii was compared with Im, and Sz was compared with Koi) (Table S1). Culture strains were maintained using plastic Petri dishes (90 × 20 mm) and PESI medium (Tatewaki, 1966) at 15 and 10°C in long day (16h:8h, light:dark) and short day (8h:16h, light:dark) conditions with 30–50 μmol m^-2^s^-1^ photon flux density.

To examine fertilization rate of gametes from parthenogenetic populations, they were crossed with male gametes from sexual populations. The crossing was performed following Hoshino et al. (2019). We added in large excess male gametes to female gametes in a small drop that settled in microscopic field and observed what proportion of the female gametes formed zygotes in five minutes. As control, crossing between female and male gametes from sexual populations was also performed. For the statistical analysis of the fertilization rate (the number of fertilized gametes and unfertilized gametes), a generalized linear mixed model (GLMM) was adopted with a binomial distribution. Models with and without sexuality (Amazon, sexual female, and sexual male) were established for the fertilization rate (identity of culture strains was considered as the random effect in both models), and the AIC values and parameters for each were examined (Table S3). The modeling and model selection were conducted in R (R Core Team 2022) using packages lme4 (Bates *et al*., 2015).

Brown algae sex pheromones are hydrophobic, cycloaliphatic unsaturated hydrocarbons, comprising eight to eleven carbon atoms (Maier and Müller, 1986). Sexual pheromone was detected using a gas chromatography/mass spectrometry (GC-MS), as in (Hoshino & Kogame, 2019). Fertile gametophytes were kept in 300 mL of sterilized seawater in a 500 mL flask at 15°C and gametes were released in the flask. Volatile secretions were trapped on Mono Trap RCC18 (GL science, Tokyo, Japan) using a closed-loop-stripping system at 15°C. After looping 12 h, absorbed volatile compounds were eluted with 50 μL of CH_2_Cl_2_ and immediately analyzed by GC-MS, using Zebron ZB-wax columns (Phenomenex, Germany) 30 m × 0.25 μm; He as the carrier gas; program rate: 45-200°C at a rate of 5°C/min. Compounds were identified using the NIST MS library and similarity search program(Daniel Siderius, 2017). Females and males from sexual populations were also analyzed as positive and negative controls respectively. Sex pheromone was also detected by olfaction in blind tests.

Several minutes after gametes release, gametes lose their motility and settle to the substratum and then change their shape from pear-shaped to spherical. We took images of spherical gametes with a Nikon Digital Sight DS-Fi1 (Nikon Corporation, Tokyo) and measured their diameter using ImageJ software (Schneider *et al*., 2012). Sixteen to 50 gametes were measured for each individual strain. For the statistical analysis of the gamete size, a GLMM was adopted with a normal distribution.

To examine parthenogenetic development, gametes were cultivated at 15°C in long day conditions as described above. We recoded the number of germinated gametes and non-germinated gametes after 24h cultivation and the number of cells of each germling after five-day cultivation. For each individual, between 40–160 gametes/germlings were scored. For the statistical analysis of the germination rate and cell numbers of germlings, a GLMM was adopted with a binominal distribution and a Poisson distribution, respectively (Table S3). All statistical analysis were performed in R.

### Genome sequencing and assembly

Genomes of *S. lomentaria* (strain Kn2f) and *S. promiscuus* (strain Ii3ax and Im6f) were sequenced to be used as references for analyses of RNA-seq data. Prior the genome DNA extraction, the gametophytes were cultivated with antibiotics (Penicillin G 50 mg/L, Ampicillin 50 mg/L, Chloramphenicol 5 mg/L, and Kanamycin 0.5g/L) for one week to remove eventual bacterial contamination. The gametophytes were frozen in liquid nitrogen and crushed to powder using TissueLyser II (QIAGEN) and genomic DNA was extracted using OmniPrep^TM^ for Plant (GBiosciences) as in (Ahmed *et al*., 2014; Cossard *et al*., 2022). The libraries of Kn2f and Im6f were sequenced on an illumina Nextseq 2000 with paired-end reads of 150 bp. *S. promiscuus* strain Ii3ax was sequenced using Oxford Nanopore Sequencing (ONT) MinION. OmniPrepTM and NucleoBond for high molecular weight DNA were used to extract the genomic DNA. ONT sequencing was performed on a R9.4.1 flow cell with the SQK-LSK110 ligation sequencing Kit.

The *S. promiscuus* nanopore ONT reads were basecalled by the ONT basecaller Guppy version 6.3.8+d9e0f64 and the configuration file dna_r9.4.1_450bps_sup.cfg. The fastq reads were filtered for bacterial contamination by using a conjunction of Kraken2 version 2.1.2 (Wood *et al*., 2019) and blastn version 2.9.0+ (Altschul *et al*., 1990) in combination with the NCBI nt database (download date 2022-07-01).The filtered reads were assembled by SMARTdenovo (options J 500, k 32) (Liu *et al*., 2021). Super scaffolding was performed by ONT medaka_consensus version 1.7.2 (https://github.com/nanoporetech/medaka) and default options. Gap closing was performed by PBJelly version v15.8.24 (https://github.com/esrice/PBJelly) (English *et al*., 2012) using *S. promiscuus* strain Im6f Illumina read contigs.

For *S. lomentaria,* the quality of the scaffolded genomes was assessed using QUAST v.5.0.2 (Mikheenko *et al*., 2018), and statistics are shown in Table S1.

### Genome structural annotation

All genomes were soft-masked using Repeatmasker v 4.1.2 after building a *de-novo* transposable elements and repeats database with RepeatModeler v2.0.3 (Flynn *et al*., 2020). Two runs of BRAKER2 v2.1.6 (Brůna *et al*., 2021) were used to predict gene sets used for all downstream analyses: (i) using input predicted protein from model species *Ectocarpus* sp (EctsiV2_prot_LATEST.tfa, (Cock *et al*., 2010; Cormier *et al*., 2017) and chordaria linearis (Ectocarpales, Cossard et al., 2022) and (ii) using RNAseq data from the same individual sequenced mapped onto the genome with tophat2 (v2.1.1; Kim et al., 2013). TSEBRA (Gabriel *et al*., 2021) was then used to select transcripts from the previous two runs of BRAKER2 with parameters recommended to include ab-initio predicted genes. Genome annotation completeness was assessed by BUSCO v3 (Waterhouse *et al*., 2018) against the eukaryote and stramenopile gene set Odb10 (Manni et al., 2021).

### RNA sequencing

Gametophytes were cultivated in plastic Petri dishes (90 × 20 mm) and sterile natural sea water from the North Sea enriched with full strength PESI medium at 10°C in long day (16h:8h, light:dark) conditions with LED lighting of 20 μmol m^-2^s^-1^ photon flux density. Medium was renewed every week until gamete collection.

RNA libraries from at least three replicate individuals per sex for each lineage (Table S1) were prepared from gametes using NEBNext® Single Cell/Low Input RNA Library Prep Kit for Illumina (New England Biolabs). Mature gametophytes release gametes just after the medium renewal. Gametes were collected by phototaxis: *Scytosiphon* gametes have negative phototaxis, so freshly released gametes accumulate on the opposite side of a light source in a Petri dish. Between 300–10,500 gametes in 0.5 μl of medium were transferred to a PCR tube and incubated for five minutes in darkness at 10°C to allow gametes to settle on the surface of the tube. After incubation, 5 μl of the NEBNext Cell Lysis Buffer was added to the tube, and then proceeded immediately to reverse transcription and library preparation, following the manufacture’s protocol. Libraries were sequenced on an illumina Nextseq 2000 with paired-end reads of 150 bp.

### Phylogenetic tree construction based on RNA-seq data

To examine phylogenetic relationships among parthenogenetic and sexual lineages, phylogenetic trees were generated based on assembled RNA-seq data. Adapter sequences and low-quality reads were removed using Trimmomatic (Bolger *et al*., 2014). The filtered reads were mapped on either *S. lomentaria* or *S. promiscuus* genome using Tophat2 v.2.1.1. The resulted bam files were sorted and used for genome-guided transcriptome assembly by Trinity v.2.13.2 (Grabherr *et al*., 2011). Coding sequences (CDS) were predicted for the longest isoform of each assembled transcriptome using Transdecoder v5.7.0. To remove transcriptomes from potential contamination, amino acid sequences from the predicted CDS were blasted against algae_peptide database (algal sequences downloaded from the NCBI and the JGI databases) using DIAMOND (Buchfink *et al*., 2015) with sensitive mode, and proteins without blast-hit were removed. To remove highly identical amino acid sequences, CD-HIT (Li & Godzik, 2006; Fu *et al*. 2012) was used with the parameter ‘*-c 0.95*’. Then, using the filtered datasets, single copy orthologues among the samples were predicted by OrthoFinder v.2.5.4 (Emms & Kelly, 2015). The 78 single copy orthologues were detected, and the nucleotide sequences of 53 genes whose missing rate was < 20% were concatenated and used for a phylogenetic analysis. The phylogenetic analysis was performed by IQ-TREE v2.1.4 (Minh *et al*., 2020) with the flag ‘-MFP+MERGE’ with 1000 of ultrafast bootstrapping.

### Divergence time estimation

The divergence time between lineages was estimated using cox1, cox3, and rbcL. We constructed a time-calibrated tree of 23 species of Ectocarpales and Laminariales (Table S15) based on the mitochondrial *cox*1, *cox*3 and chloroplast *rbc*L (total 2819 bp) using BEAST version 2.7.4 (Drummond *et al*., 2005; Bouckaert *et al*., 2019) with the following settings: substitution models of GTR+Γ for *rbc*L and *cox*3, and HKY+Γ for *cox*1, an optimized relaxed clock, Yule Model prior, and 131,815,000 generations of Markov chain Monte Carlo (MCMC) with sampling every 1,000 generations. In addition, we specified a prior on the root age of *Nereocystis luetkeana* and *Pelagophycus porra* as a calibration point like (Silberfeld *et al*., 2010). We assumed that the two species are monophyletic and the lower boundary of their divergence time is 13 Ma (exponential distribution with mean and offset of 2.0 and 13.0, respectively). Stationarity of the MCMC ran was checked by Tracer version 1.7.1 (Rambaut *et al*., 2018). A maximum clade credibility tree with median node heights was constructed with TreeAnnotator (Drummond *et al*., 2006) with burn-in of 10%. The constructed tree was visualized by FigTree version 1.4.3. Note that we could not estimate the divergence times of a subset of the samples because there were not enough polymorphisms: in *S. promiscuus*, sexual and Amazon populations (Koi and Sz, and Im and Ii) have identical cox1 haplotypes, so divergence time cannot be estimated. Regarding *S. lomentaria*, we cannot distinguish sexual from asexuals using cox1 (Hoshino *et al*., 2021). The divergence times between *S. lomentaria* and *S. promiscuus* was estimated as 13.7 Ma (median, [95%HPD: 8.1–21.9 Ma]), and between *S. promiscuus* Koi and *S. promiscuus* Im populations as 0.76 Ma (median, [95%HPD: 0.1–1.8 Ma]).

### Expression quantification and identification of sex-biased genes in sexual populations

RNAseq reads were removed if the leading or trailing base had a Phred score <3, or if the sliding window Phred score, averaged over four bases, was <15. Reads shorter than 36 bases were discarded (as well as pair of reads, if one of the pair was <36 bases long). Trimmomatic-processed RNAseq reads from each library were used to quantify gene expression with kallisto v 0.46.2 (Bray *et al*., 2016) using 31 bp-long kmers and predicted transcript of each species. RNAseq libraries were all composed of paired-end reads. A gene was considered expressed in a given species and/or a given sex when at least two-third of samples displayed an expression level above 0.4 counts. Estimates of sex-biased gene expression in dioicous sexual populations were obtained using read count matrices as input for the DESeq2 package (Love *et al*., 2014) in R 3.6.3. P-values were corrected for multiple testing using Benjamini and Hochberg’s algorithm in DESeq2, applying an adjusted P-value cut-off of 0.05 for differential expression analysis. Genes with a minimum of 2-fold change expression level between sexes were retained as sex-biased. We plotted heatmap of expression levels in log2(TPM + 1) for each sample within each species using the ComplexHeatmap R package (Gu, 2022).

### Gene expression at U-specific genes in females versus Amazons

Expression values (transcript per million, TPM) of U-SDR gens in males, females, and Amazon populations of *S. lomentaria* and *S. promiscuus* were compared in the context of sex linkage genes. The *Scytosiphon* sex linked genes were extracted from(Lipinska *et al*., 2017) based on orthology with *Ectocarpus* genes. If an *Ectocarpus* gene model had hits on multiple *Scytosiphon* gene models (*Scytosiphon* gene models were split or truncated), the *Scytosiphon* gene models were merged to represent a complete *Ectocarpus* model. To identify significant differences at the gene expression, a Wilcoxon test was performed for each U-sex linked gene.

### Orthology and evolutionary sequence divergence between *Scytosiphon species*

We inferred single-copy orthologs (SCO) between the two species using Orthofinder (v2.5.2) with default parameters (Emms & Kelly, 2015). We used kallisto v 0.46.2 to quantify expression levels of SCO. Orthologous proteins between species pairs were aligned with MAFFT v7.453(Katoh *et al*., 2009) the alignments were curated with Gblocks v0.91b (Talavera & Castresana, 2007) and back-translated to nucleotides using translatorX (Abascal *et al*., 2010). We used these nucleotide alignments as input for phylogenetic analysis by maximum likelihood (PAML4, CodeML) (Yang, 2007) to infer pairwise dN/dS (ω) with F3×4 model of codon frequencies. We retained orthologs with 0 < dS < 2 as valid for further analysis. We compared species evolutionary rates separately for female-biased, male-biased, and unbiased genes with Mann-Whitney ranked test. No orthogroup presented an inconsistent sex-bias across the two species, i.e., biased in one sex in one species and in the opposite sex in the other species.

### Polymorphism (SNPs) inference within Scytosiphon species

RNA-seq data of all strains shown in Figure 1C were passed on to detect synonymous and non-synonymous mutations in the context of amazon specific mutations (female versus Amazon). *S. promiscuus and S. lomentaria* data were treated the same way, but kept separate by the two strains. Filtered RNA-seq data from all samples were used to detect variants according to the pipeline by (Haas *et al*., 2020). Software updates were accomplished for GATK version 4.2.6.1 (McKenna *et al*., 2010)and gmap-gsnap version 2021-12-17 (Wu & Nacu, 2010). Only variants which passed the filter steps (read depth >=9 and supporting reads >=7) and located on exon features were kept for further analysis.

The reference genome of *S. lomentaria* is a female (Kn2f) and the reference genome of *S. promiscuus* is an Amazon (Ii3ax). The filter was set to a read coverage of >=9 reads and >=7 reads have to support the alternative base (according to Haas et al., 2020). Non-synonymous variants were detected using SNPEff version 5.1d (Cingolani *et al*., 2012) run in default mode. Exclusive variants represent the unique section of the intersection between female, male and Amazon samples. Candidates are variant positions present only in Amazon samples and all Amazon strains have to contain the variant. Since the variant calling may miss some SNPs or InDels, each of the *S. promiscuus* candidate positions were manually checked with JBrowse (if the mapping files [bam] do show exclusive ax SNPs). Twenty-five candidate positions were identified as Amazon-specific and used for the further analyses. The 25 positions are located on 16 gene models. 15 gene models have orthologs on *S. lomentaria*. By manually checking the variant positions on the 15 *S. lomentaria* orthologs, one single specific Amazon variant was identified, across both species. The *S. lomentaria* gene is anno1.g11893 (pos. Contig91_RagTag:1006619), the *S. promiscuus* gene is anno1.g1469 (pos. Contig91:1065707).

### Estimation of pN/pS

Variants were filtered to retain SNP present in at least two individuals per group, with a minimum coverage of 7 reads and a minimum average phred quality of 20. SNP variants were annotated using snpEff with custom database for *S promiscuus* and *S lomentaria* (Cingolani *et al*., 2012). We used these annotated variants to compute the ratio of non-synonymous to synonymous polymorphism per site (pN/pS = π) per gene, following (Nei & Gojobori, 1986) using a custom script.

To test for differences between sexual systems we used generalized linear models, with binomial distribution, to analyse the proportion of genes that displayed SNPs and the proportion of variable sites in genes that had SNPs. We included the log-transformed expression level (in TPM) as independent variable and the gene identity as a random factor. Models were computed separately for each *Scytosiphon* species. We used linear mixed models to infer the effect of the sexual system on π separately per species. These analyses were repeated on the SCO gene subset.

### Expression changes between sexual and asexual females

We compared the expression levels of female-and male-biased genes in sexual versus asexual females with Mann-Whitney ranked tests to investigate the possible masculinization and defeminization of Amazon expression profiles, separately for each species.

Convergent changes associated with transitions to asexuality were investigated on single-copy orthologs (SCO) inferred across the two species. We use DESeq2 including expression data from sexual and asexual (Amazon) females only (males were removed) to model the gene expression as a function of the species, the sexual system (sexual or asexual) and the interaction between these two variables. We retained convergently differential expressed genes performing likelihood ratio test when they were significantly affected by sexual system but not species nor by the interaction of species by sexual system.

The SuperExactTest package in R was used to perform exact multi-set intersection tests in order to determine whether representation of sex-biased genes among genes showing convergent expression gene was greater than expected by chance (p-value < 0.05).

To further characterize the role and fate of sex-biased genes during the transition to asexuality we used a similar approach to (Cossard *et al*., 2022). We compared the expression profiles of expressed genes (in log2(TPM + 1)) between sexual and Amazon females by computing the Pearson correlation coefficients of expression levels, within species, between the mean expression across all sexual females and each asexual female sample, separately for female-biased, male-biased, and non-biased genes. We compared Pearson coefficients of regression within each species, using the cocor package (Diedenhofen & Musch, 2015), considering gene expression profiles of SBG and unbiased genes within sexes as dependent. We report the P-value based on (Meng *et al*., 1992; Diedenhofen & Musch, 2015).

Evolutionary sequence divergence (d_N_/d_S_) of single-copy orthologs with convergent expression shifts was compared to the rest of SCO to infer whether these genes experienced different selective pressure (Mann-Whitney ranked tests).

### Functional annotation analysis

Predicted genes for each species were blasted against the NCBI non-redundant (nr) protein database with Diamond (v2.0.15)(Buchfink *et al*., 2015). Functional annotation was performed using BLAST2GO ((Conesa & Götz, 2008), as well as the InterProScan (v5.59-91.0) prediction of putative conserved protein domains (Quevillon *et al*., 2005). Gene set enrichment analysis (GSEA) was carried out separately for each species and each geneset (female-biased, male-biased) as well as for genes with convergent expression shift with asexuality (combining GO-terms inferred for each species for genes within orhtogroups) using Fisher’s exact test implemented in the TopGO package, with the weight01 algorithm (Alexa & Rahnenfuhrer, 2020). We investigated enrichment in terms of molecular function ontology and reported significant GO-terms with P-value < 0.05. All statistical analyses were performed in R 4.2.3, plots were produced with ggplot2 in R (Wickham, 2016).

### Structure prediction

Primary sequences were aligned with ClustalW and T-Coffee, followed by JPred secondary structure prediction through Jalview (Li, 2003; Cole *et al*., 2008; Waterhouse *et al*., 2009) to determine the putative domain boundaries. A custom script to predict the protein models with AlphaFold2 was utilized to generate all protein models in this analysis (Baltzis *et al*., 2022). Only protein models with the highest ranked predicted local distance difference test (pLDDT) score were selected.

## Acknowledgements

This work was supported by the MPG, the ERC (grant n. 864038 to SMC), the JSPS Overseas Research Fellowships (to MH), the BMBF-funded de.NBI Cloud within the German Network for Bioinformatics Infrastructure (de.NBI) (031A532B, 031A533A, 031A533B, 031A534A, 031A535A, 031A537A, 031A537B, 031A537C, 031A537D, 031A538A). SMC is supported by the Moore Foundation (GBMF11489) and the Bettencourt-Schuller Foundation. We thank Masanori Hiraoka for help with sampling and phenotyping *S. promiscuus*, Elena Avdievich for help with Nanopore sequencing of *S. promiscuus*, Denis Roze and Sylvain Glemin for helpful discussions, and Carol Featherstone for assistance in the preparation of the manuscript. Vikram Alva and John R. Weir shared the custom AlphaFold2 script that was utilized to generate the protein models.

## Supplemental Figures Legends

**Figure S1.**
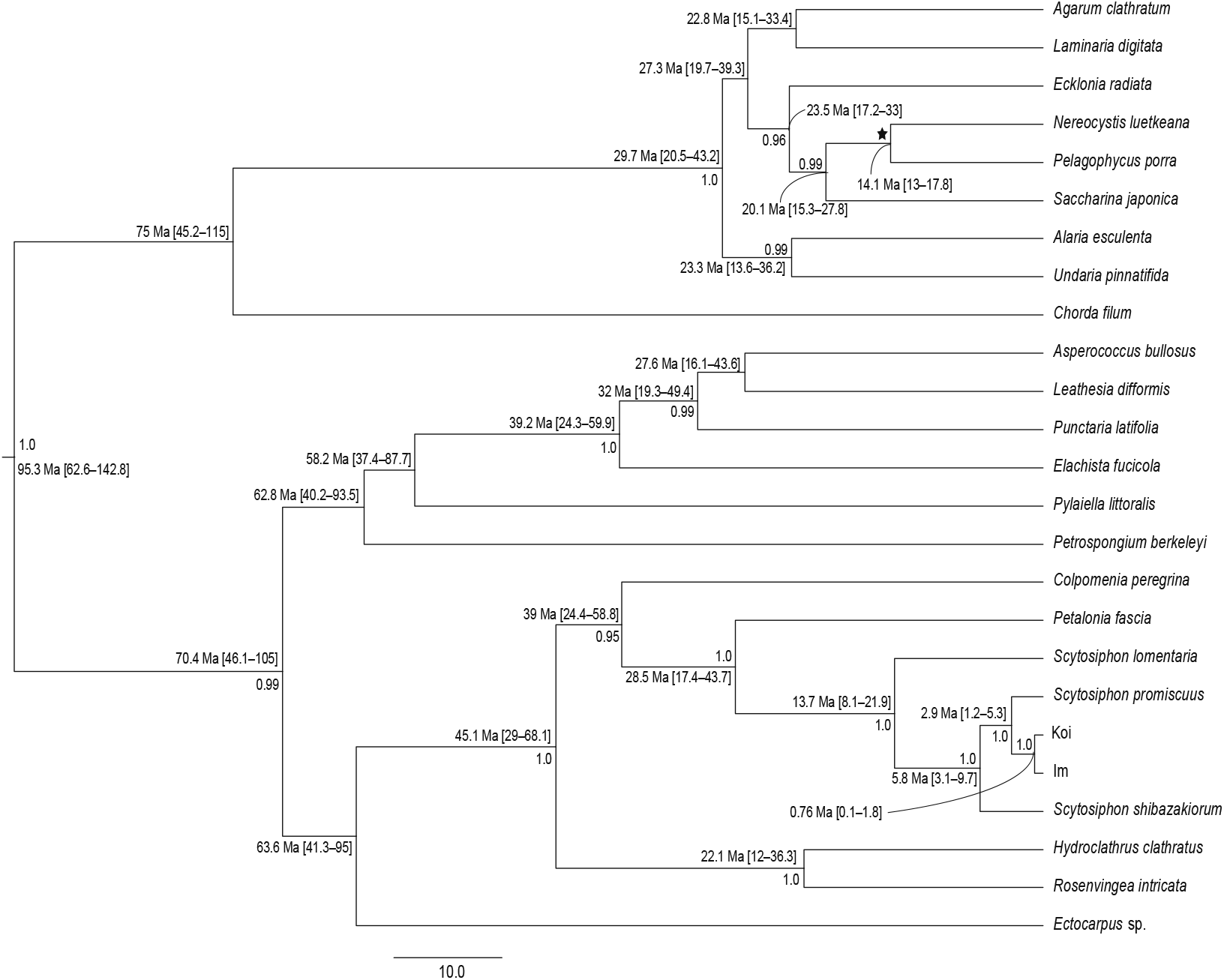
Phylogenetic tree showing divergence time. Posterior probability of > 0.95 and the median value of estimated divergence time (95% highest probability density in square brackets) are given to each node. The black star indicates the node used as a calibration point, an exponential prior with 13 Ma as lower boundary.

**Figure S2.**
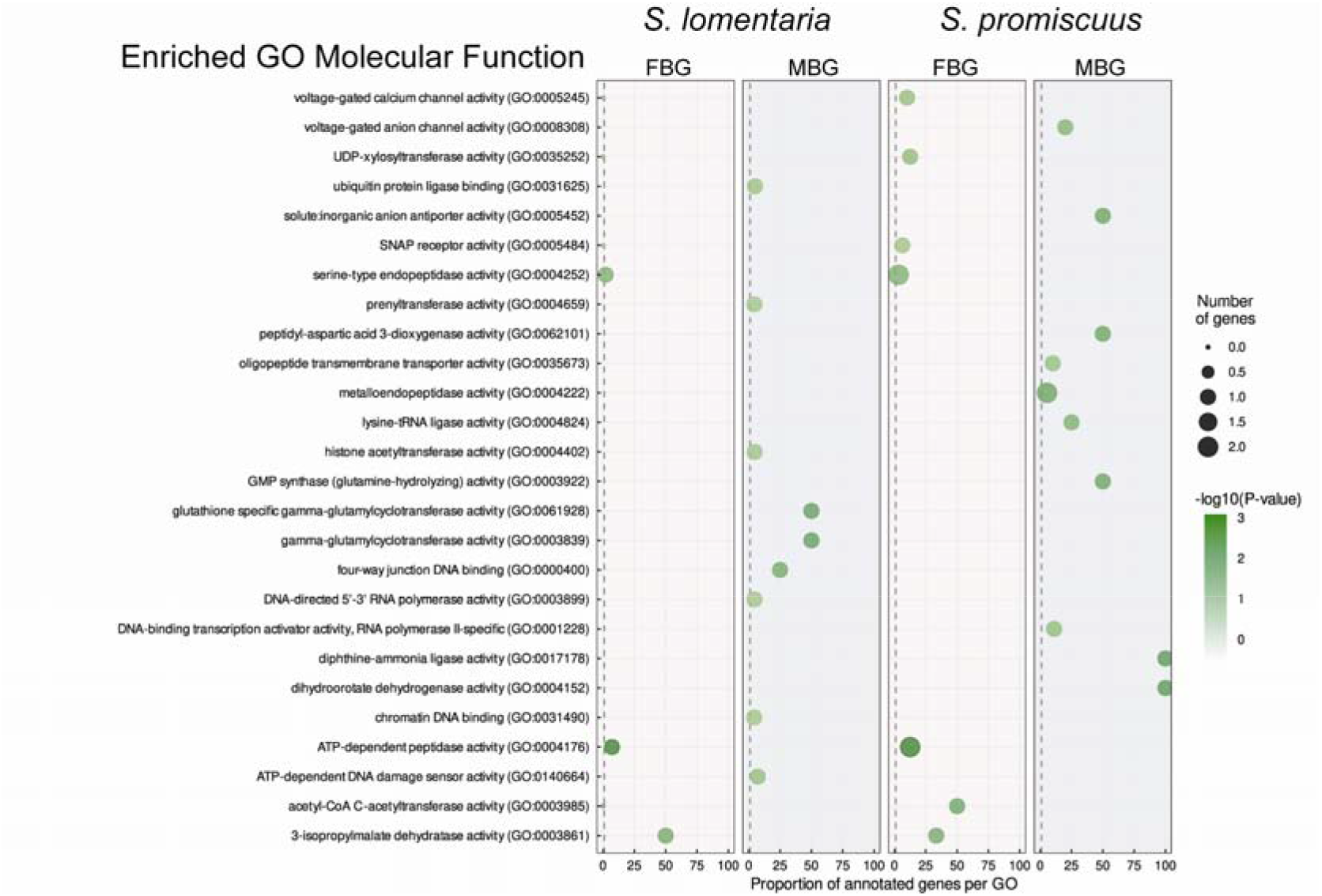
GO term enrichment (Molecular function) in male-and female-biased genes in *S. lomentaria* and *S. promiscuus*.

**Figure S3.**
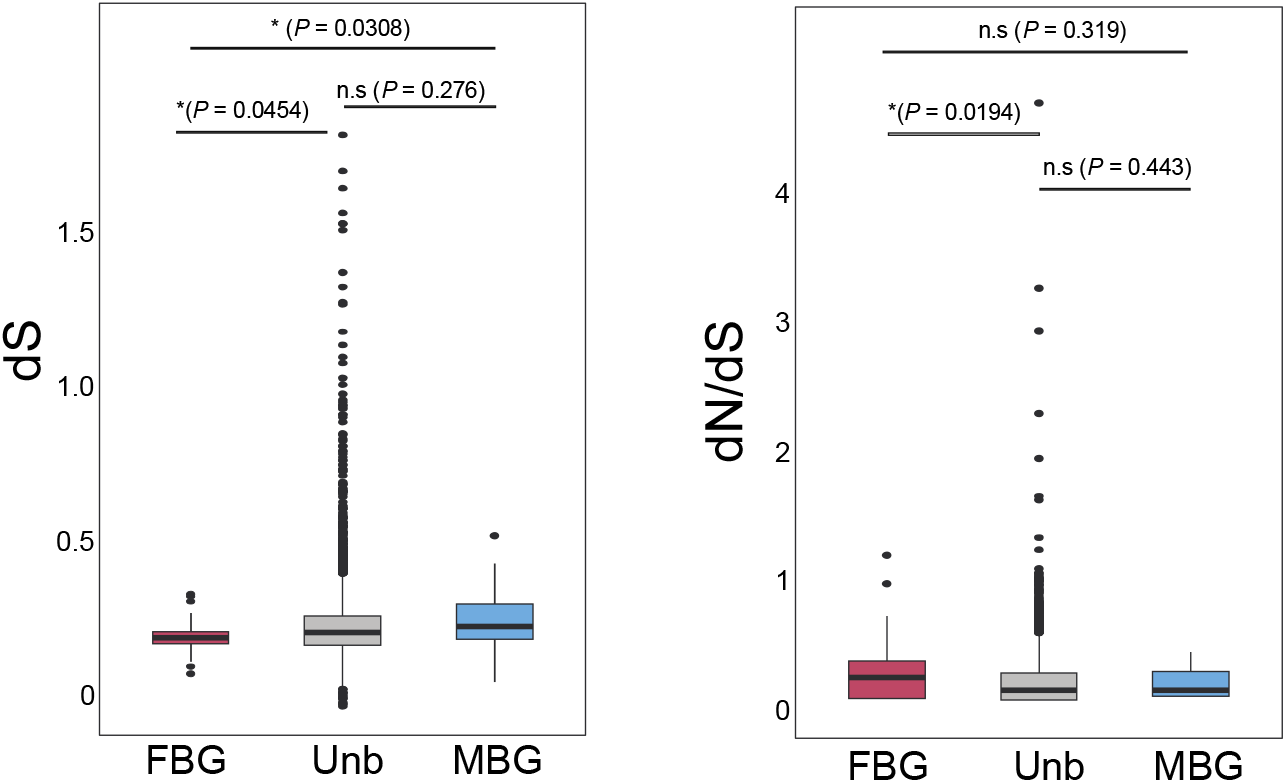
Evolutionary rates measured as dN/dS between species pairs (*S. lomentaria*/*S.promiscuus*) for unbiased, female-biased (FBG), and male-biased genes (MBG). The statistical tests are permutation Mann-Whitney ranked tests; the *p-*values are displayed in parentheses.

**Figure S4.**
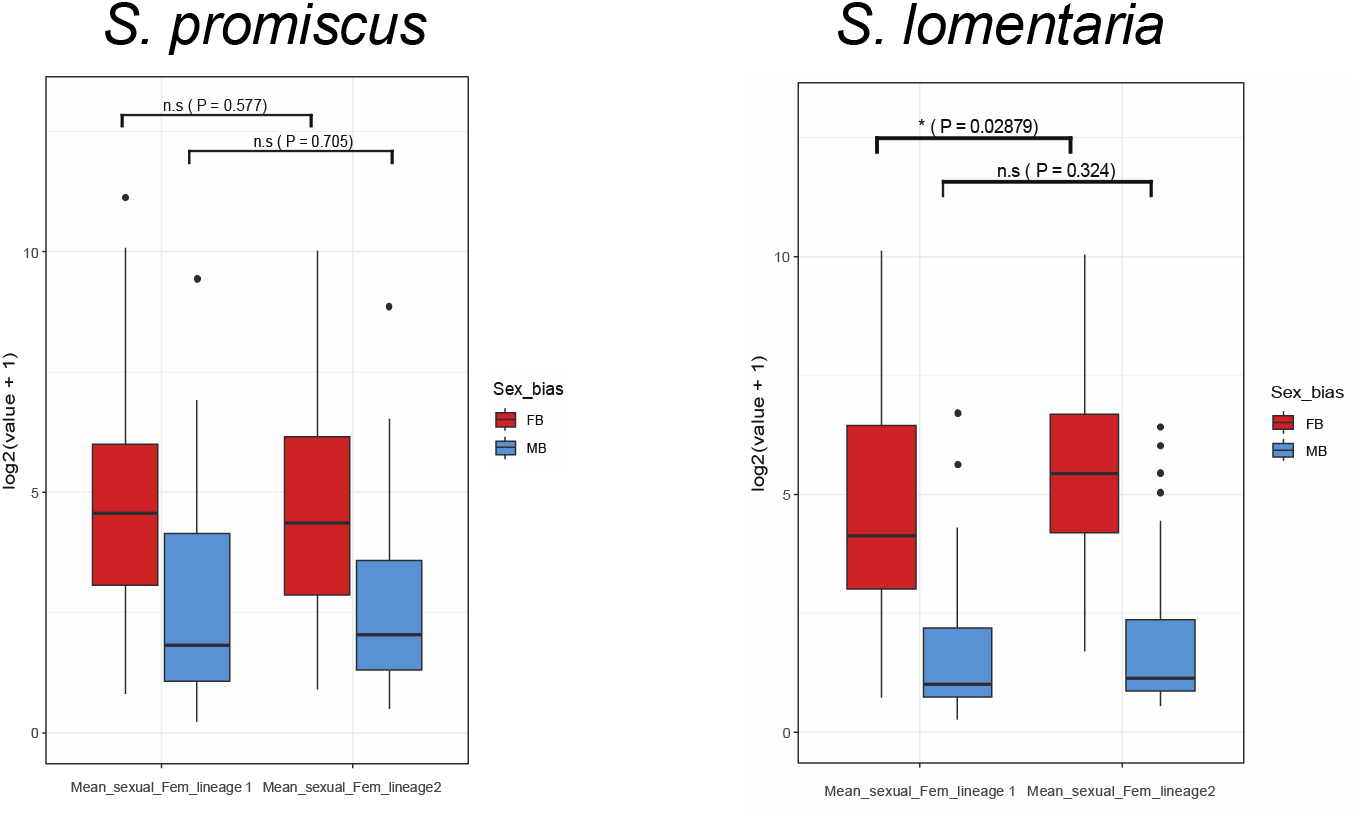
Changes in sex-biased gene expression when two sexual lineages are compared.

**Figure S5.**
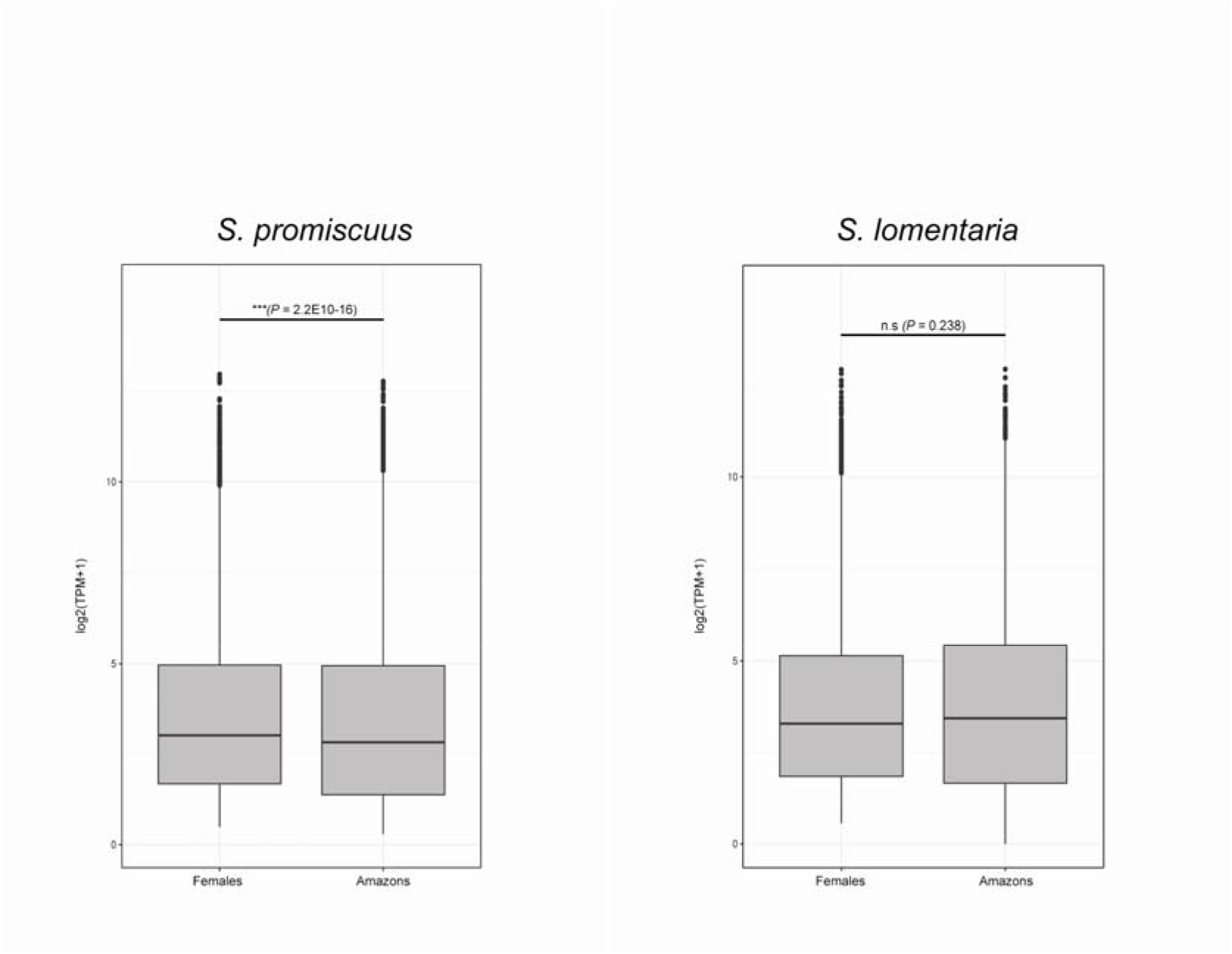
Comparison of unbiased gene expression levels in sexual and asexual populations, in log2(TPM+1). Boxes represent the interquartile range (25th and 75th percentiles) of the data, the line inside the box represents the median, whiskers represent the largest/smallest value within 1.5 times interquartile range above and below the 75th and 25th percentile, respectively. Statistical tests are Mann-Whitney ranked tests. Number of analysed genes are presented inside brackets.

**Figure S6.**
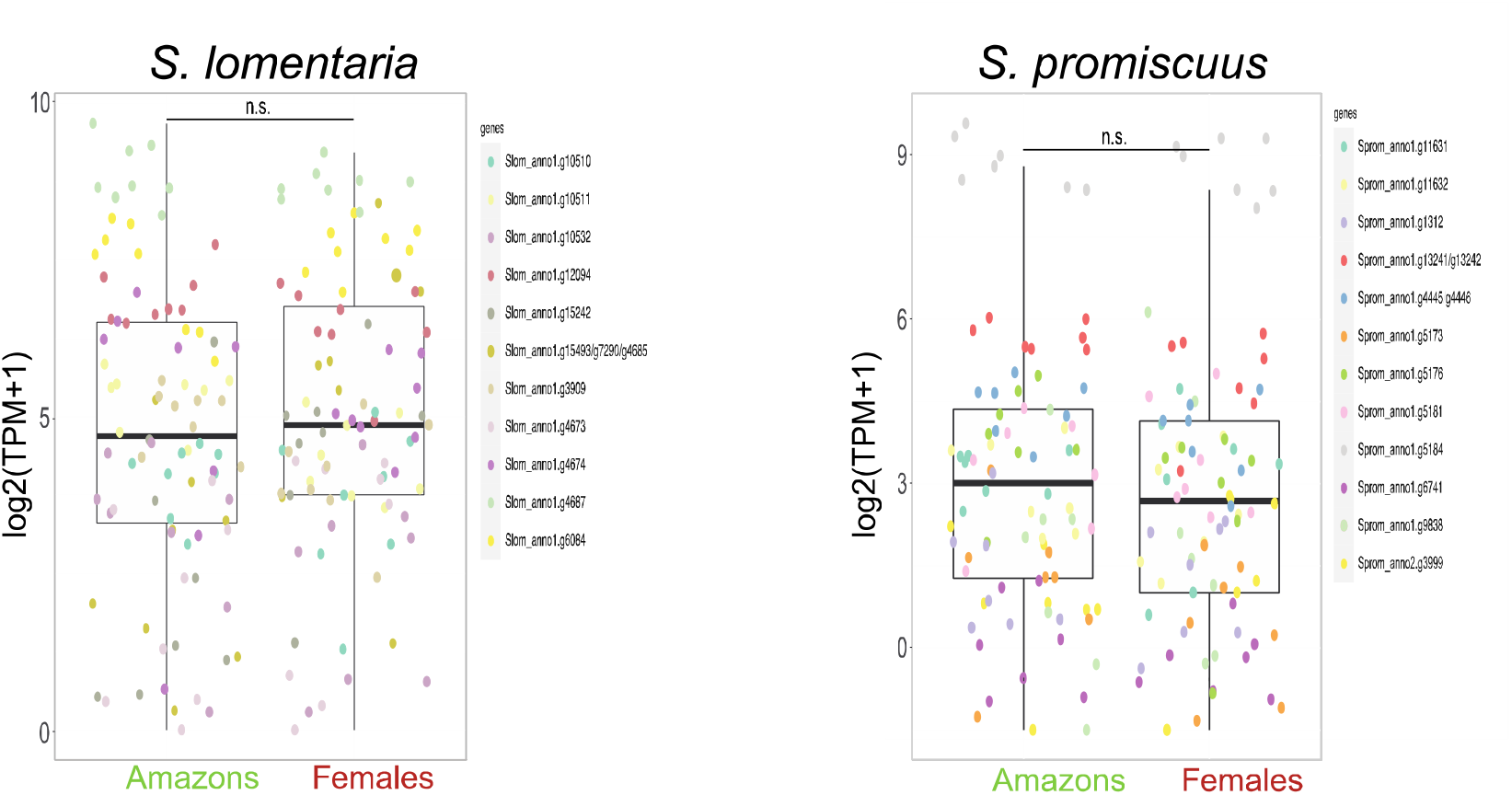
Mean expression level of U-linked genes in Amazon populations compared to ancestral sexual populations.

## Supplemental Tables Legends

**Table S1**. Strains used in this study.

**Table S2**. Identification of the sex of each individual gametophte from natural populations of S. promiscuus.

**Table S3**. Tables summarizing the phenotypes in male, females and asexuals in the different populations of Scytosiphon. Fertilisation success was scored in each of the populations by counting proportion of zygotes. Pheromone was measured either by olfactation in a blind test or by GC-MS. Attachment of male flagellum was measured by microscope observation (blind test). Gamete fusion was scored 24h after mixing male and female gametes by observation of zygote production (confirmed with presence of two eyespots), as in Coelho et al. 2011.

**Table S4**. GLMM models to test significance of differences in phenotypes of sexual and asexual females

**Table S5**. Genome sequencing details and structural annotation.

**Table S6**. Number of expressed genes in *S. lomentaria and S. promiscuus* lineages and differential expression (DE) levels.

**Table S7**. Functional annotation (GO terms) and expression level (in trancript per million, TPM) of *S. promiscuus* genes.

**Table S8**. Functional annotation (GO terms) and expression level (in trancript per million, TPM) of *S. lomentaria* genes.

**Table S9**. Significantly enriched Go-term (Molecular function Gene Ontology) per geneset of sex-biased genes (Fisher’s exact tests).

**Table S10**. Statistics of likelyhood ratio tests (LRT) for sexual system for all single-copy orthologs between *S. promiscuus* and *S. lomentaria*. ‘Convergent’ refers to genes whose expression change during transition to asexuality is consistent in both species

**Table S11**. Expression (transcript per million, TPM) of U-SDR gens in males, females and aseuxal populations of *S. lomentaria* (Slom) and *S. promiscuus* (Sprom). Sex linkage was from Lipinska et al. 2017 based on orthology with *Ectocarpus* genes that are U-linked and PCR validation in *S. lomentaria*. Note that we assume that these genes are also sex linked in *S. promiscuus* given that they are also sex linked in other brown algae (Lipinska et al. 2017). Statistically different expression (Wilcoxon tests) is shown for the comparison females versus asexual populations.

**Table S12**. P-values (Mann-Whitney ranked tests) for evolutionary rates of single-copy orthologs. Statistical significance is indicated in bold.

**Table S13**. Significantly enriched Gene Ontology (GO)-terms of OGs with convergent expression associated with the shift to asexuality, in comparison to all Orthogroups (Fisher’s exact tests).

**Table S14**. Statistics of the RNA-seq variant calling, separated by total, female (f), amazon (ax) and male (m).

**Table S15**. Accession numbers of the datasets used for construction of the phylogenetic tree to estimate times of divergence between *Scytosiphon* populations and species

## References

1. Abascal F, Zardoya R, Telford MJ. 2010. TranslatorX: multiple alignment of nucleotide sequences guided by amino acid translations. Nucleic acids research 38: W7–13.

2. Abbott JK. 2011. Intra-locus sexual conflict and sexually antagonistic genetic variation in hermaphroditic animals. Proceedings. Biological sciences 278: 161–169.

3. Ahmed S, Cock JM, Pessia E, Luthringer R, Cormier A, Robuchon M, Sterck L, Peters AF, Dittami SM, Corre E, et al. 2014. A haploid system of sex determination in the brown alga Ectocarpus sp. Current biology: CB 24: 1945–1957.

4. Alexa A, Rahnenfuhrer J. 2020. Enrichment Analysis for Gene Ontology.

5. Altschul SF, Gish W, Miller W, Myers EW, Lipman DJ. 1990. Basic local alignment search tool. Journal of Molecular Biology 215: 403–410.

6. Arun A, Peters NT, Scornet D, Peters AF, Cock JM, Coelho SM. 2013. Non-cell autonomous regulation of life cycle transitions in the model brown alga Ectocarpus. The New Phytologist 197: 503–510.

7. Baltzis A, Mansouri L, Jin S, Langer BE, Erb I, Notredame C. 2022. Highly significant improvement of protein sequence alignments with AlphaFold2 (PL Martelli, Ed.). Bioinformatics 38: 5007–5011.

8. Bast J, Parker DJ, Dumas Z, Jalvingh KM, Tran Van P, Jaron KS, Figuet E, Brandt A, Galtier N, Schwander T. 2018. Consequences of Asexuality in Natural Populations: Insights from Stick Insects (S Wright, Ed.). Molecular Biology and Evolution 35: 1668– 1677.

9. Bates D, Mächler M, Bolker B, Walker S. 2015. Fitting Linear Mixed-Effects Models Using lme4. Journal of Statistical Software 67.

10. Bolger AM, Lohse M, Usadel B. 2014. Trimmomatic: a flexible trimmer for Illumina sequence data. Bioinformatics (Oxford, England) 30: 2114–2120.

11. Bothwell JH, Marie D, Peters AF, Cock JM, Coelho SM. 2010a. Role of endoreduplication and apomeiosis during parthenogenetic reproduction in the model brown alga Ectocarpus. The New phytologist 188: 111–121.

12. Bothwell JH, Marie D, Peters AF, Cock JM, Coelho SM. 2010b. Cell cycles and endocycles in the model brown seaweed, Ectocarpus siliculosus. Plant Signaling & Behavior 5: 1473–1475.

13. Bouckaert R, Vaughan TG, Barido-Sottani J, Duchêne S, Fourment M, Gavryushkina A, Heled J, Jones G, Kühnert D, De Maio N, et al. 2019. BEAST 2.5: An advanced software platform for Bayesian evolutionary analysis (M Pertea, Ed.). PLOS Computational Biology 15: e1006650.

14. Brandt A, Schaefer I, Glanz J, Schwander T, Maraun M, Scheu S, Bast J. 2017. Effective purifying selection in ancient asexual oribatid mites. Nature Communications 8: 873.

15. Bray NL, Pimentel H, Melsted P, Pachter L. 2016. Near-optimal probabilistic RNA-seq quantification. Nature biotechnology 34: 525–527.

16. Brůna T, Hoff KJ, Lomsadze A, Stanke M, Borodovsky M. 2021. BRAKER2: automatic eukaryotic genome annotation with GeneMark-EP+ and AUGUSTUS supported by a protein database. NAR genomics and bioinformatics 3: lqaa108.

17. Buchfink B, Xie C, Huson DH. 2015. Fast and sensitive protein alignment using DIAMOND. Nature methods 12: 59–60.

18. Cingolani P, Platts A, Wang LL, Coon M, Nguyen T, Wang L, Land SJ, Lu X, Ruden DM. 2012. A program for annotating and predicting the effects of single nucleotide polymorphisms, SnpEff: SNPs in the genome of Drosophila melanogaster strain w1118D; iso-2; iso-3. Fly 6: 80–92.

19. Cock JM, Sterck L, Rouzé P, Scornet D, Allen AE, Amoutzias G, Anthouard V, Artiguenave F, Aury J-M, Badger JH, et al. 2010. The Ectocarpus genome and the independent evolution of multicellularity in brown algae. Nature 465: 617–621.

20. Coelho SMB, Brownlee C, Bothwell JHF. 2008. A tip-high, Ca(2+) -interdependent, reactive oxygen species gradient is associated with polarized growth in Fucus serratus zygotes. Planta 227: 1037–1046.

21. Coelho SM, Cock JM. 2020. Brown Algal Model Organisms. Annual Review of Genetics 54: 71–92.

22. Coelho SM, Godfroy O, Arun A, Le Corguillé G, Peters AF, Cock JM. 2011. OUROBOROS is a master regulator of the gametophyte to sporophyte life cycle transition in the brown alga Ectocarpus. Proceedings of the National Academy of Sciences of the United States of America 108: 11518–11523.

23. Coelho SM, Taylor AR, Ryan KP, Sousa-Pinto I, Brown MT, Brownlee C. 2002. Spatiotemporal Patterning of Reactive Oxygen Production and Ca2+ Wave Propagation in Fucus Rhizoid Cells. The Plant Cell 14: 2369–2381.

24. Coelho SM, Umen J. 2021. Switching it up: algal insights into sexual transitions. Plant reproduction 34: 287–296.

25. Cole C, Barber JD, Barton GJ. 2008. The Jpred 3 secondary structure prediction server. Nucleic Acids Research 36: W197–W201.

26. Conesa A, Götz S. 2008. Blast2GO: A comprehensive suite for functional analysis in plant genomics. International Journal of Plant Genomics 2008: 619832.

27. Connallon T, Knowles LL. 2005. Intergenomic conflict revealed by patterns of sex-biased gene expression. Trends in genetics._: TIG 21: 495–499.

28. Cormier A, Avia K, Sterck L, Derrien T, Wucher V, Andres G, Monsoor M, Godfroy O, Lipinska A, Perrineau M-M, et al. 2017. Re-annotation, improved large-scale assembly and establishment of a catalogue of noncoding loci for the genome of the model brown alga Ectocarpus. The New phytologist 214: 219–232.

29. Cossard GG, Godfroy O, Nehr Z, Cruaud C, Cock JM, Lipinska AP, Coelho SM. 2022. Selection drives convergent gene expression changes during transitions to co-sexuality in haploid sexual systems. Nature ecology & evolution 6: 579–589.

30. Daniel Siderius. 2017. NIST Standard Reference Simulation Website - SRD 173.

31. Dedryver C-A, Le Gallic J-F, Mahéo F, Simon J-C, Dedryver F. 2013. The genetics of obligate parthenogenesis in an aphid species and its consequences for the maintenance of alternative reproductive modes. Heredity 110: 39–45.

32. Diedenhofen B, Musch J. 2015. cocor: a comprehensive solution for the statistical comparison of correlations. PloS one 10: e0121945.

33. Drummond DA, Bloom JD, Adami C, Wilke CO, Arnold FH. 2005. Why highly expressed proteins evolve slowly. Proceedings of the National Academy of Sciences of the United States of America 102: 14338–14343.

34. Drummond AJ, Ho SYW, Phillips MJ, Rambaut A. 2006. Relaxed Phylogenetics and Dating with Confidence (D Penny, Ed.). PLoS Biology 4: e88.

35. Emms DM, Kelly S. 2015. OrthoFinder: solving fundamental biases in whole genome comparisons dramatically improves orthogroup inference accuracy. Genome biology 16: 157.

36. English AC, Richards S, Han Y, Wang M, Vee V, Qu J, Qin X, Muzny DM, Reid JG, Worley KC, et al. 2012. Mind the Gap: Upgrading Genomes with Pacific Biosciences RS Long-Read Sequencing Technology (Z Liu, Ed.). PLoS ONE 7: e47768.

37. Felsenstein J. 1971. INBREEDING AND VARIANCE EFFECTIVE NUMBERS IN POPULATIONS WITH OVERLAPPING GENERATIONS. Genetics 68: 581–597.

38. Flynn JM, Hubley R, Goubert C, Rosen J, Clark AG, Feschotte C, Smit AF. 2020. RepeatModeler2 for automated genomic discovery of transposable element families. Proceedings of the National Academy of Sciences of the United States of America 117: 9451–9457.

39. Fu G, Kinoshita N, Nagasato C, Motomura T. 2014. Fertilization of Brown Algae: Flagellar Function in Phototaxis and Chemotaxis. In: Sawada H, Inoue N, Iwano M, eds. Sexual Reproduction in Animals and Plants. Tokyo: Springer Japan, 359–367.

40. Gabriel L, Hoff KJ, Brůna T, Borodovsky M, Stanke M. 2021. TSEBRA: transcript selector for BRAKER. BMC Bioinformatics 22: 566.

41. Glemin S, Galtier N. 2012. Genome evolution in outcrossing versus selfing versus asexual species. Methods in molecular biology (Clifton, N.J.) 855: 311–335.

42. Grabherr MG, Haas BJ, Yassour M, Levin JZ, Thompson DA, Amit I, Adiconis X, Fan L, Raychowdhury R, Zeng Q, et al. 2011. Full-length transcriptome assembly from RNA-Seq data without a reference genome. Nature Biotechnology 29: 644.

43. Haas FB, Fernandez-Pozo N, Meyberg R, Perroud P-F, Göttig M, Stingl N, Saint-Marcoux D, Langdale JA, Rensing SA. 2020. Single Nucleotide Polymorphism Charting of P. patens Reveals Accumulation of Somatic Mutations During in vitro Culture on the Scale of Natural Variation by Selfing. Frontiers in Plant Science 11: 813.

44. Hartfield M. 2016. Evolutionary genetic consequences of facultative sex and outcrossing. Journal of Evolutionary Biology 29: 5–22.

45. Heesch S, Serrano-Serrano M, Barrera-Redondo J, Luthringer R, Peters AF, Destombe C, Cock JM, Valero M, Roze D, Salamin N, et al. 2021. Evolution of life cycles and reproductive traits: insights from the brown algae. Journal of Evolutionary Biology n/a.

46. Hojsgaard D, Hörandl E. 2019. The Rise of Apomixis in Natural Plant Populations. Frontiers in Plant Science 10: 358.

47. Hollister JD, Greiner S, Wang W, Wang J, Zhang Y, Wong GK-S, Wright SI, Johnson MTJ. 2015. Recurrent Loss of Sex Is Associated with Accumulation of Deleterious Mutations in Oenothera. Molecular Biology and Evolution 32: 896–905.

48. Hoshino M, Hiruta SF, Croce ME, Kamiya M, Jomori T, Wakimoto T, Kogame K. 2021. Geographical parthenogenesis in the brown alga Scytosiphon lomentaria (Scytosiphonaceae): Sexuals in warm waters and parthenogens in cold waters. Molecular Ecology 30: 5814–5830.

49. Hoshino M, Kogame K. 2019. Parthenogenesis is rare in the reproduction of a sexual field population of the isogamous brown alga Scytosiphon (Scytosiphonaceae, Ectocarpales) (M Cock, Ed.). Journal of Phycology 55: 466–472.

50. Hoshino M, Okino T, Kogame K. 2018. Parthenogenetic female populations in the brown alga Scytosiphon lomentaria (Scytosiphonaceae, Ectocarpales): decay of a sexual trait and acquisition of asexual traits. Journal of phycology.

51. Hoshino M, Okino T, Kogame K. 2019. Parthenogenetic female populations in the brown alga Scytosiphon lomentaria (Scytosiphonaceae, Ectocarpales): decay of a sexual trait and acquisition of asexual traits (M Cock, Ed.). Journal of Phycology 55: 204–213.

52. Huylmans AK, Macon A, Hontoria F, Vicoso B. 2021. Transitions to asexuality and evolution of gene expression in Artemia brine shrimp. Proceedings of the Royal Society B: Biological Sciences 288: 20211720.

53. Johnston SE, Gratten J, Berenos C, Pilkington JG, Clutton-Brock TH, Pemberton JM, Slate J. 2013. Life history trade-offs at a single locus maintain sexually selected genetic variation. Nature 502: 93–95.

54. Katoh K, Asimenos G, Toh H. 2009. Multiple alignment of DNA sequences with MAFFT. Methods in molecular biology (Clifton, N.J.) 537: 39–64.

55. Kearney M. 2005. Hybridization, glaciation and geographical parthenogenesis. Trends in Ecology & Evolution 20: 495–502.

56. Keightley PD, Otto SP. 2006. Interference among deleterious mutations favours sex and recombination in finite populations. Nature 443: 89–92.

57. Kitayama T, Kawai H, Yoshida T. 1992. Dominance of female gametophytes in field populations of Cutleria cylindrica (Cutleriales, Phaeophyceae) in the Tsugaru Strait, Japan. Phycologia 31: 449–461.

58. Li K-B. 2003. ClustalW-MPI: ClustalW analysis using distributed and parallel computing. Bioinformatics 19: 1585–1586.

59. Lipinska, Ahmed S, Peters AF, Faugeron S, Cock JM, Coelho SM. 2015a. Development of PCRDbased markers to determine the sex of kelps (C Wicker-Thomas, Ed.). PLoS ONE 10: e0140535.

60. Lipinska, Cormier A, Luthringer R, Peters AF, Corre E, Gachon CMM, Cock JM, Coelho SM. 2015b. Sexual dimorphism and the evolution of sex-biased gene expression in the brown alga Ectocarpus. Molecular biology and evolution 32: 1581– 1597.

61. Lipinska AP, Serrano-Serrano ML, Cormier A, Peters AF, Kogame K, Cock JM, Coelho SM. 2019. Rapid turnover of life-cycle-related genes in the brown algae. Genome biology 20: 35.

62. Lipinska AP, Toda NRT, Heesch S, Peters AF, Cock JM, Coelho SM. 2017. Multiple gene movements into and out of haploid sex chromosomes. Genome Biology 18: 104.

63. Liu H, Wu S, Li A, Ruan J. 2021. SMARTdenovo: a de novo assembler using long noisy reads. Gigabyte 2021: 1–9.

64. Liu L-J, Zheng H-Y, Jiang F, Guo W, Zhou S-T. 2014. Comparative Transcriptional Analysis of Asexual and Sexual Morphs Reveals Possible Mechanisms in Reproductive Polyphenism of the Cotton Aphid (KY Zhu, Ed.). PLoS ONE 9: e99506.

65. Love MI, Huber W, Anders S. 2014. Moderated estimation of fold change and dispersion for RNA-seq data with DESeq2. Genome biology 15: 550.

66. Luthringer R, Cormier A, Peters AF, Cock JM, Coelho SM. 2015. Sexual dimorphism in the brown algae. 1: 11–25.

67. Ma PWK, Roelofs WL. 1995. Calcium involvement in the stimulation of sex pheromone production by PBAN in the European corn borer, Ostrinia nubilalis (Lepidoptera: Pyralidae). Insect Biochemistry and Molecular Biology 25: 467–473.

68. Maier I. 1995. Brown algal pheromones. 11.

69. Mank JE. 2017. The transcriptional architecture of phenotypic dimorphism. Nature Ecology & Evolution 1: 0006.

70. McKenna A, Hanna M, Banks E, Sivachenko A, Cibulskis K, Kernytsky A, Garimella K, Altshuler D, Gabriel S, Daly M, et al. 2010. The Genome Analysis Toolkit: a MapReduce framework for analyzing next-generation DNA sequencing data. Genome Research 20: 1297–1303.

71. Meng X, Rosenthal R, Rubin DB. 1992. Comparing correlated correlation coefficients. Psychological Bulletin 111: 172–175.

72. Mignerot L, Avia K, Luthringer R, Lipinska AP, Peters AF, Cock JM, Coelho SM. 2019. A key role for sex chromosomes in the regulation of parthenogenesis in the brown alga Ectocarpus. PLoS genetics 15: e1008211.

73. Mignerot L, Coelho SM. 2016. The origin and evolution of the sexes: Novel insights from a distant eukaryotic linage. Comptes rendus biologies 339: 252–257.

74. Mikheenko A, Prjibelski A, Saveliev V, Antipov D, Gurevich A. 2018. Versatile genome assembly evaluation with QUAST-LG. Bioinformatics 34: i142–i150.

75. Minh BQ, Schmidt HA, Chernomor O, Schrempf D, Woodhams MD, Von Haeseler A, Lanfear R. 2020. IQ-TREE 2: New Models and Efficient Methods for Phylogenetic Inference in the Genomic Era (E Teeling, Ed.). Molecular Biology and Evolution 37: 1530–1534.

76. Nei M, Gojobori T. 1986. Simple methods for estimating the numbers of synonymous and nonsynonymous nucleotide substitutions. Molecular Biology and Evolution.

77. Neiman M, Hehman G, Miller JT, Logsdon JM, Taylor DR. 2010. Accelerated Mutation Accumulation in Asexual Lineages of a Freshwater Snail. Molecular Biology and Evolution 27: 954–963.

78. Neiman M, Sharbel TF, Schwander T. 2014. Genetic causes of transitions from sexual reproduction to asexuality in plants and animals. Journal of evolutionary biology 27: 1346–1359.

79. Normark BB, Moran NA. 2000. Testing for the accumulation of deleterious mutations in asexual eukaryote genomes using molecular sequences. Journal of Natural History 34: 1719–1729.

80. Parker DJ, Bast J, Jalvingh K, Dumas Z, Robinson-Rechavi M, Schwander T. 2019. Sex-biased gene expression is repeatedly masculinized in asexual females. Nature communications 10: 4638.

81. Peters AF, Scornet D, Ratin M, Charrier B, Monnier A, Merrien Y, Corre E, Coelho SM, Cock JM. 2008. Life-cycle-generation-specific developmental processes are modified in the immediate upright mutant of the brown alga Ectocarpus siliculosus. Development 135: 1503–1512.

82. Quevillon E, Silventoinen V, Pillai S, Harte N, Mulder N, Apweiler R, Lopez R. 2005. InterProScan: protein domains identifier. Nucleic acids research 33: W116–120.

83. Rambaut A, Drummond AJ, Xie D, Baele G, Suchard MA. 2018. Posterior Summarization in Bayesian Phylogenetics Using Tracer 1.7 (E Susko, Ed.). Systematic Biology 67: 901–904.

84. Schneider CA, Rasband WS, Eliceiri KW. 2012. NIH Image to ImageJ: 25 years of image analysis. Nature Methods 9: 671–675.

85. Silberfeld T, Leigh JW, Verbruggen H, Cruaud C, de Reviers B, Rousseau F. 2010. A multi-locus time-calibrated phylogeny of the brown algae (Heterokonta, Ochrophyta, Phaeophyceae): Investigating the evolutionary nature of the ‘brown algal crown radiation’. Molecular phylogenetics and evolution 56: 659–674.

86. Simon J-C, Delmotte F, Rispe C, Crease T. 2003. Phylogenetic relationships between parthenogens and their sexual relatives: the possible routes to parthenogenesis in animals: ROUTES TO PARTHENOGENESIS IN ANIMALS. Biological Journal of the Linnean Society 79: 151–163.

87. Stearns SC. 1989. Trade-Offs in Life-History Evolution. Functional Ecology 3: 259.

88. Szövényi P, Devos N, Weston DJ, Yang X, Hock Z, Shaw JA, Shimizu KK, McDaniel SF, Wagner A. 2014. Efficient Purging of Deleterious Mutations in Plants with Haploid Selfing. Genome Biology and Evolution 6: 1238–1252.

89. Talavera G, Castresana J. 2007. Improvement of phylogenies after removing divergent and ambiguously aligned blocks from protein sequence alignments. Systematic Biology 56: 564–577.

90. Tatewaki M. 1966. Formation of a Crustaceous Sporophyte with Unilocular Sporangia in Scytosiphon lomentaria. Phycologia 6: 62–66.

91. Tewari R, Bailes E, Bunting KA, Coates JC. 2010. Armadillo-repeat protein functions: questions for little creatures. Trends in Cell Biology 20: 470–481.

92. Umen J, Coelho S. 2019. Algal sex determination and the evolution of anisogamy. Annual Review of Microbiology 73: 267–291.

93. Waterhouse AM, Procter JB, Martin DMA, Clamp M, Barton GJ. 2009. Jalview Version 2—a multiple sequence alignment editor and analysis workbench. Bioinformatics 25: 1189–1191.

94. Waterhouse RM, Seppey M, Simão FA, Manni M, Ioannidis P, Klioutchnikov G, Kriventseva EV, Zdobnov EM. 2018. BUSCO Applications from Quality Assessments to Gene Prediction and Phylogenomics. Molecular Biology and Evolution 35: 543–548.

95. Wickham H. 2016. ggplot2: Elegant Graphics for Data Analysis.

96. Wood DE, Lu J, Langmead B. 2019. Improved metagenomic analysis with Kraken 2. Genome Biology 20: 257.

97. Wu TD, Nacu S. 2010. Fast and SNP-tolerant detection of complex variants and splicing in short reads. Bioinformatics 26: 873–881.

98. Yang Z. 2007. PAML 4: phylogenetic analysis by maximum likelihood. Molecular biology and evolution 24: 1586–1591.

99. Ye Z, Molinier C, Zhao C, Haag CR, Lynch M. 2019. Genetic control of male production in Daphnia pulex. Proceedings of the National Academy of Sciences 116: 15602–15609.

